# Hierarchical control of bacterial growth efficiency by substrate and taxonomy

**DOI:** 10.64898/2026.07.26.740805

**Authors:** Vaibhhav Sinha, Seppe Kuehn

## Abstract

Heterotrophic bacteria utilize organic carbon as a source of energy and material for growth. The balanced allocation of carbon between these two processes, termed growth efficiency, is a key physiological property of microbes because it links energy and biomass production, and determines the fraction of carbon lost as CO_2_. However, we do not understand what controls growth efficiency in microbes or how it correlates with taxonomy and resource identity. Here, we develop a quantitative high-throughput method for measuring CO_2_ production during bacterial growth. For 23 bacterial strains spanning three phyla, grown on two carbon substrates, we quantify growth efficiency by measuring dynamic CO_2_ production and carbon accumulation in biomass. Intra-phylum comparisons show that glycolytic substrates yield higher efficiency growth (less CO_2_ produced per biomass carbon) than gluconeogenic substrates. However, growth efficiency varies as much across resources as it does across phyla. A physiological model shows that variation in the growth efficiency depends on the ATP produced per respired CO_2_ and the ATP needed per biomass on a given substrate, suggesting phylum level differences in energy supply and demand dictate differences in growth efficiency. This theory predicts no global correlation between growth rate and growth efficiency, a finding our data support. Finally, we report phylum-level variation in the dynamics of CO_2_ production, which we link to the presence of overflow metabolism on glycolytic substrates. This study revises our understanding of how taxonomy and substrate identity impact carbon allocation during growth and sets the stage for understanding CO_2_ production in communities.

**Significance:** Bacteria are responsible for roughly half of all biological CO_2_ production on the planet, much of it in soils. How organic carbon flows through microbial metabolism sets the rate at which soils return CO_2_ to the atmosphere. Bacterial growth efficiency, defined by the fraction of consumed carbon that is retained as biomass rather than released as CO_2_, determines the fluxes of carbon into the atmosphere. At present, we do not understand what controls growth efficiency in bacteria. We develop a new quantitative measurement of growth efficiency which we apply to 23 soil bacteria from three phyla on two substrates. We show that efficiency is structured hierarchically, at the highest level substrate identity defines efficiency which then varies substantially across taxa. So both the identity of species involved and the compounds they consume determine the rate of carbon loss.

---

Heterotrophic microbes in soils consume organic carbon and respire several times more CO_2_ each year than all human activity combined (1, 2). Much of this heterotrophic respiration is bacterial. Here we seek to understand what controls the fraction of organic carbon a bacterium retains as biomass versus releases as CO_2_.

Growing bacteria utilize carbon to generate energy, biomass, and excreted organic byproducts (Fig. 1A). The bacterial growth efficiency (BGE) is the ratio of carbon in biomass to the sum of carbon in biomass and respired CO_2_. As a result, bacterial growth efficiency reflects the balance between cellular demands for energy and material. Increasing production of CO_2_ reflects greater energy production at the expense of carbon for biomass. Since nearly half of soil organic matter is comprised of dead microbial cells (3), the balance between biomass production and CO_2_ release plays a defining role in the global carbon cycle.

**Fig. 1.**
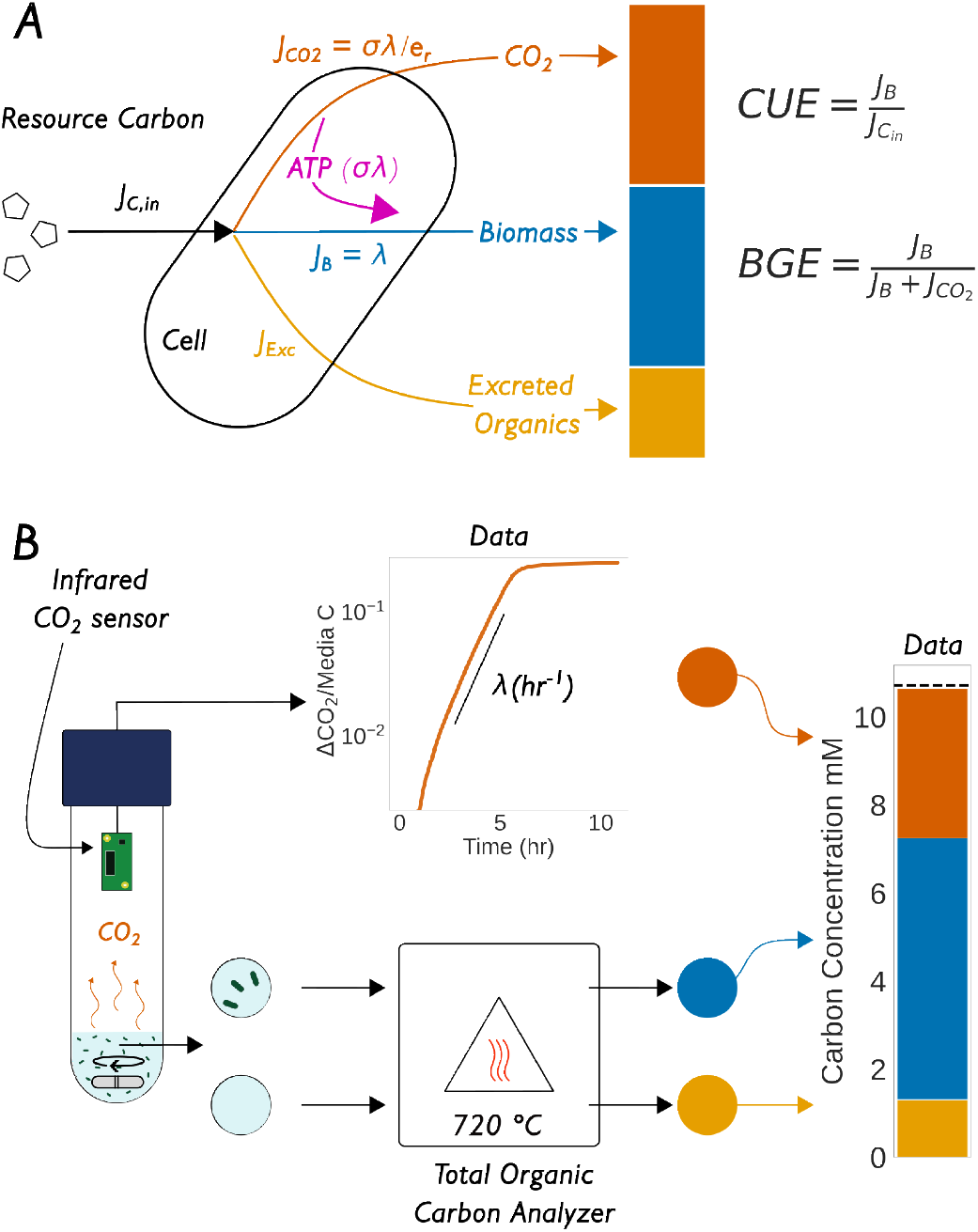
Carbon allocation metrics and quantification. **(A)** During aerobic respiration, carbon utilized (*J*_*C,in*_) by growing bacteria has three fates: biomass (*J*_*B*_), respired 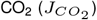 and excreted organic byproducts (*J*_exc_) as shown. Two dimensionless ratios are used to quantify this allocation: Carbon Use Efficiency (defined below), CUE = *J*_*B*_*/J*_*C*,in_, and Bacterial Growth Efficiency, 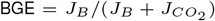. BGE and CUE are identical when *J*_exc_ = 0. **(B)** Experimental workflow: Bacteria are cultured in defined media with fixed initial organic carbon inside hermetically sealed vessels with non-destructive infrared CO_2_ sensors. The sensors report real-time headspace CO_2_ (orange curve, inset). At the end of growth, total organic carbon in biomass and supernatant, and dissolved inorganic carbon in the liquid, are quantified on a Total Organic Carbon (TOC) analyzer, yielding specific growth rate *λ* and the endpoint excreted carbon and biomass carbon.

Community-scale measurements establish that soil carbon retention varies with temperature, substrate stoi-chiometry, and nutrient limitation (4, 5). These measurements motivate ecosystem-scale models that make microbial physiology explicit (6, 7), but such models remain poorly constrained: their projections diverge widely because the physiological, chemical, and taxonomic bases of carbon allocation are unknown. This gap could be alleviated by measurements of BGE for individual organisms in defined environmental conditions.

At the organism scale, BGE has been measured for individual strains growing on a single resource (8–11). However, distinct measurement methodologies, assumptions, resources, and taxa have given rise to sometimes contradictory findings (see Discussion). For example, some studies report positive correlation between growth rate and BGE (8), while others report a negative association (9). Thus it remains unclear which aspects of cellular physiology that vary across taxa control this critical carbon allocation process.

In contrast, Andersen and von Meyenburg (12) quantify BGE for *Escherichia coli* growing on ten substrates and observe a correlation between BGE and growth rate driven by higher BGE and growth rates on glycolytic than gluconeogenic substrates. Within classes of resources (e.g. sugars or acids) they observe wide variation in growth rates at roughly constant BGE (Fig. S1). Mori and colleagues report similar findings (13).

Following Basan (14) and Mori (13), a simple model balancing energy supply and demand predicts that BGE should vary with energy yield per unit substrate and energy demand per unit biomass, but not directly on growth rate (assuming cellular maintenance energies are small, see SI Section 1, Eqs. 14-17). In this view, the energy demand is constant for a strain, and differences in BGE between substrate classes likely arise from differences in net energy yields per CO_2_ on those substrates, with wide variations in growth rate potentially arising from proteomic and not energetic constraints (15). What remains unclear is how BGE varies across distinct taxa on a given resource or across multiple resources. Here we aim to address this gap.

We measure BGE for 23 heterotrophic soil bacteria from three phyla (Pseudomonadota, Actinomycetota, Bacillota) grown on glucose or succinate. We develop custom respirometry devices that quantify headspace CO_2_ in real time, combined with direct quantification of organic and inorganic carbon at the end of growth. For every strain that grows on both resources, BGE on glucose exceeds BGE on succinate, consistent with the higher ATP yield per CO_2_ from glycolytic catabolism. However, BGE varies strongly with taxonomy: Actinomycetota are systematically more efficient than Pseudomonadota and Bacillota on both substrates. In contrast to prior studies, BGE and growth rate are uncorrelated within resources. TCA cycle and electron-transport-chain gene content statistically predict taxonomic variation in BGE, consistent with the idea that energy yield per unit substrate and its correlation with CO_2_ production is one factor that reflects differences in BGE. Therefore, the fate of carbon during aerobic respiration is determined both by substrate identity and taxonomy in a manner that does not strongly correlate with growth rate but is predictable from genomic content.

## Results

### The definition of growth efficiency

Two ratios are widely used in the literature to quantify carbon allocation during heterotrophic growth (Fig. 1A). Carbon use efficiency (CUE) is the fraction of carbon taken up that is retained in biomass and is sometimes called yield. Bacterial growth efficiency (BGE) is the ratio of carbon allocated to biomass to carbon in biomass plus carbon lost as CO_2_. These ratios can be understood by considering mass balance of fluxes during exponential growth 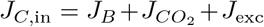 (Fig. 1A). From this,

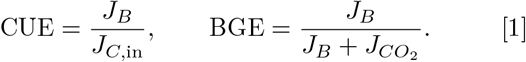

CUE and BGE are equal when *J*_exc_ = 0 and otherwise differ by the fraction of carbon lost to excretion. In what follows, we focus on BGE for populations performing aerobic respiration, and we identify conditions when *J*_*exc*_ *>* 0 and discuss this further below.

During balanced growth, the demand for energy to produce biomass is given by *J*_*E*_ = *σλ* where *σ* is the ATP cost per unit biomass carbon and *λ* is the growth rate (*J*_*B*_ = *λ*). This must balance energy production which for respiratory metabolism is proportional to the CO_2_ produced 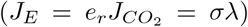. *e*_*r*_ denotes the net quantity of ATP produced per CO_2_, accounting for ATP costs of biomass precursor synthesis which varies depending on catabolic pathway utilized. Simply put, *e*_*r*_ is the net income of energy per respired CO_2_ and *σ* is the net energy cost paid per synthesized biomass carbon. Combining this energy balance with Eq. 1 gives

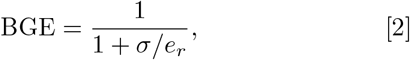

See SI Appendix, Section 1 for details and Table S1 for a summary of reported BGE or CUE values as well as the various definitions of these quantities. This result shows that BGE declines as the demand for ATP to generate biomass (*σ*) rises, and increases as the net energy produced per CO_2_ released (*e*_*r*_) rises. A similar result holds for CUE. Note there is no explicit dependence here on growth rate, so any correlations between BGE or CUE and growth rate must arise either from a dependence of growth rate on net energy yield per CO_2_ on a substrate (*e*_*r*_), changes in the energy demand per unit biomass (*σ*) as a function of growth rate, or because maintenance energies are significant relative to energetic costs of biomass. We note that *σ* consists of two components: (i) the ATP costs of generating biomass from synthesized precursors via polymerization (protein synthesis) which we expect to be roughly constant across strains and resources, and (ii) a growth-associated ATP costs beyond polymerization component (13). While the polymerization component is well understood and determined, the same cannot be said for the growth associated costs. The parameter *e*_*r*_ includes ATP from substrate level phosphorylation, electron transport, and the net ATP gain or loss from anabolism (see SI Section 1).

### A new method to quantify BGE

To enable simultaneous culturing and carbon flux quantification, we developed custom culturing devices (Fig. 1B). Each hermetically sealed culture device harbored an infrared CO_2_ sensor that enabled real-time, non-destructive measurement of CO_2_ accumulation during growth. The devices housed small (12 mL, M9 minimal medium) cultures with a large headspace (∼ 170 mL) to maintain aerobic culture conditions during growth. The dynamics of CO_2_ in time reflect phases of growth, (Fig. 1B, orange line) enabling us to infer specific growth rates (*λ*). We find that growth rates inferred from CO_2_ dynamics are consistent with those measured independently via absorbance of optical density (Fig. S9). CO_2_ production rates drop abruptly as expected when the primary carbon source is exhausted.

At the end of the growth phase, we used a destructive total carbon (TC, Shimadzu TOC-L) assay to quantify (i) total organic carbon in the biomass, (ii) total organic carbon in the liquid (spent media), (iii) total dissolved inorganic carbon (CO_2_ and bicarbonate) in the spent media. These measurements enable us to account for the fate of all the initially supplied carbon: biomass, excreted or unused organic carbon, and CO_2_.

Thus, our experiment measures an endpoint ratio: 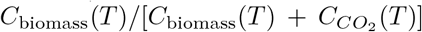, evaluated once growth has ceased (time *T*). Under balanced exponential growth with *J*_*exc*_ = 0, this endpoint ratio is identical to BGE Eq. 1 (See SI, section 2).

### BGE is hierarchically structured by substrate and taxonomy

We quantified carbon fluxes for 23 soil bacterial heterotrophs from three phyla, grown on M9 minimal medium (See SI, section 3) with either glucose or succinate as the sole carbon source. The 23 were drawn from a library of 62 isolates chosen after a screen for growth in M9 (Methods). Fig. 2A reports BGE across all strains and substrates for growth on glucose (circles) and succinate (triangles), with strains ranked by their BGE on glucose.

**Fig. 2.**
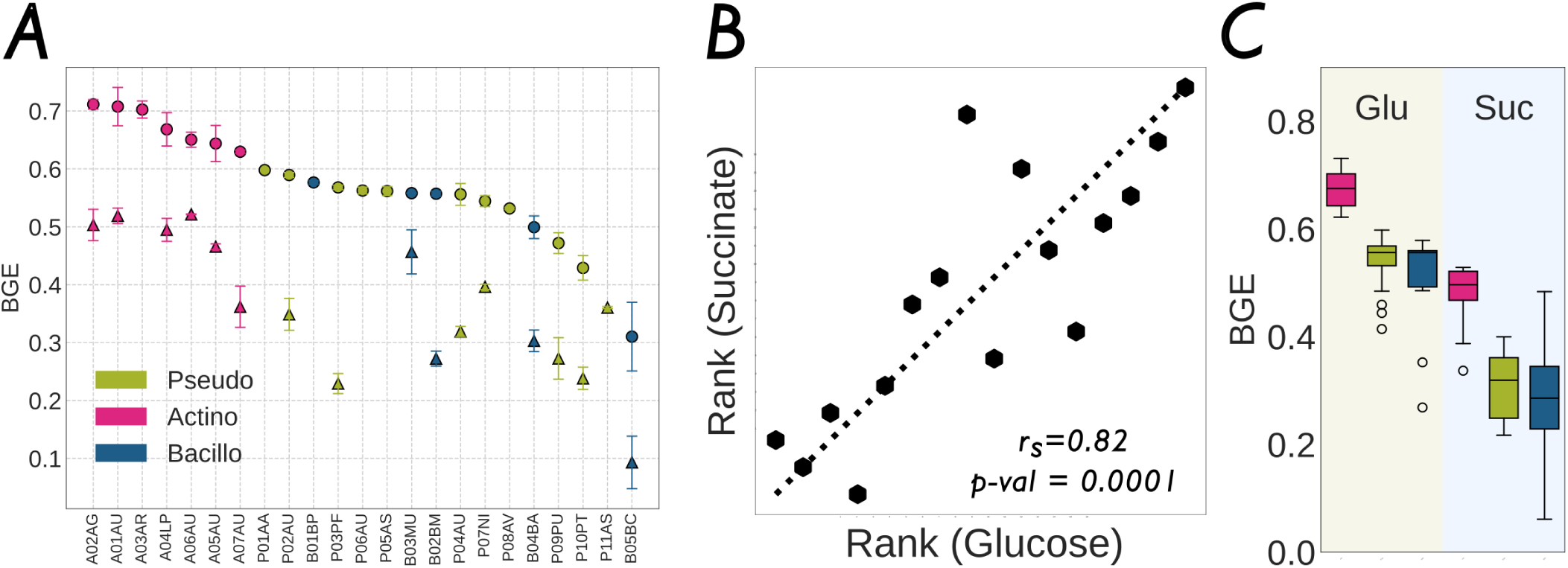
BGE is hierarchically structured by substrate and taxonomy. **(A)** BGE for each strain, ranked by BGE on glucose. Circles: glucose. Triangles: succinate. Two biological replicates per strain per resource (hollow markers). Mean BGE value for each pair of replicates is plotted using solid markers. Individual replicates as smaller, hollow markers. For every strain that grows on both substrates, BGE on glucose exceeds BGE on succinate. **(B)** Rank of BGE on succinate versus rank on glucose for the strains that grow on both substrates. The rank ordering is conserved (Spearman *r*_*s*_ = 0.82, *p* = 10^-4^), indicating that BGE reflects a strain-level physiological property that is preserved across resources. **(C)** BGE by phylum on glucose (left) and succinate (right).

From biochemical considerations we expect higher BGE from glucose than succinate due to higher *e*_*r*_ for glucose. Assuming a complete carbon oxidation, we estimate upper bounds for *e*_*r*_ ≈ 5.3 mol ATP/mol CO_2_ for glucose (glycolysis, TCA cycle, oxidative phosphorylation) and *e*_*r*_ ≈ 4.1 mol ATP/mol CO_2_ for succinate (TCA cycle entry, oxidative phosphorylation) (13, 14). See SI Section 1 for the derivation of these values. These values should be considered coarse estimates, as energy yields per CO_2_ for a given substrate are known to vary depending on the details of the metabolic pathways used(16). Nonetheless, if we assume *σ* is constant for a given strain, we expect that BGE on glucose exceeds that on succinate.

As expected from this analysis, BGE on glucose systematically exceeds BGE on succinate, for every strain that grows on both resources (Fig. 2A, circles above triangles). However, within each substrate, Actinomycetota exhibit the highest BGE, followed by Pseudomonadota and Bacillota (Fig. 2C). The magnitude of the taxonomic effect is large enough that the overall BGE of Actinomycetota on succinate approaches the overall BGE of Bacillota on glucose. Thus, taxonomy structures variation in BGE on a scale similar to variation across substrates.

Further, the strain-level rank ordering is conserved across substrates. Spearman’s rank correlation between BGE on glucose and BGE on succinate is *r*_*s*_ = 0.82 (Fig. 2B; *p* = 10^−4^). So while there is substantial taxonomic variation in growth efficiency, strain-level efficiencies exhibit conservation across resources.

BGE can be estimated from independent measurements of *σ* and *e*_*r*_ for a given strain. We have not made such measurements here, but we can use published values for model organisms to estimate plausible values for BGE. Using our upper bound estimates of *e*_*r*_ and published values of *σ* for *E. coli* (*σ* ≈ 1.2 mol ATP/mol biomass C (13)) we predict that BGE ≈ 0.81 on glucose and ≈ 0.77 on succinate. These values are higher than we observe across diverse taxa in this study (Fig. 2), due to variation in both *σ* and *e*_*r*_. We note that our measurements cannot reveal whether variation in *σ* or *e*_*r*_ is responsible for the variation in BGE across taxa on a resource. In Fig. S2 we explore different BGE values for biologically plausible ranges of *σ* and *e*_*r*_.

### BGE is not correlated with growth rate across taxa

Previous studies have reported a correlation between the growth rates of different taxa and BGE or CUE (8, 9). However, many of these studies suffer from technical challenges discussed below. So we wanted to test whether there was a correlation between BGE and growth rate in the taxa measured here.

To begin, we considered correlations across taxa on the same resource. On succinate, BGE and *λ* are uncorrelated or poorly correlated across all strains (Fig. 3A; Pearson *r* = -0.16, p-value = 0.82.). Correlations across strains intra-phylum are also not significant (Table 1). This lack of correlation indirectly supports the simple physiological model above. We assume that growth rate does not depend strongly on *e*_*r*_, so we expect no correlation. This assumption has not previously been tested, but is supported in studies on *E. coli* during aerobic growth (12, 13). Thus, our data on succinate support the claim that for a given resource, there is little correlation between growth efficiency and growth rate during aerobic growth.

**Table 1.** Pearson correlations between BGE and *λ*. Top: by phylum and carbon source. Second: by phylum and overflow status on glucose. Third: pooled across phyla by resource condition. Bottom: By phyla (inter-resource). p-values were computed from a one-sided permutation test (1000 shuffles) against the null of no positive BGE-*λ* correlation.

| | | Pearson $r$ | $p$-value |
| --- | --- | --- | --- |
| *By carbon source and phylum* |  |  |  |
| Succinate | Pseudomonadota | 0.05 | 0.43 |
|  | Actinomycetota | 0.23 | 0.23 |
|  | Bacillota | 0.18 | 0.32 |
| *By phylum and overflow status on glucose* |  |  |  |
| Overflow | Pseudomonadota | 0.88 | 0.01 |
|  | Actinomycetota | — | — |
|  | Bacillota | 0.21 | 0.44 |
| No overflow | Pseudomonadota | 0.62 | 0.29 |
|  | Actinomycetota | −0.08 | 0.57 |
|  | Bacillota* | * small sample size |  |
| *By resource condition (all phyla pooled)* |  |  |  |
| Succinate |  | −0.16 | 0.82 |
| Glucose (no overflow) |  | 0.083 | 0.42 |
| Glucose (overflow) |  | 0.76 | 0.01 |
| Overall |  | 0.07 | 0.25 |
| *By phyla (inter-resource)* |  |  |  |
| Pseudomonadota |  | 0.21 | 0.2 |
| Actinomycetota |  | 0.52 | 0.03 |
| Bacillota |  | 0.55 | 0.04 |

**Fig. 3.**
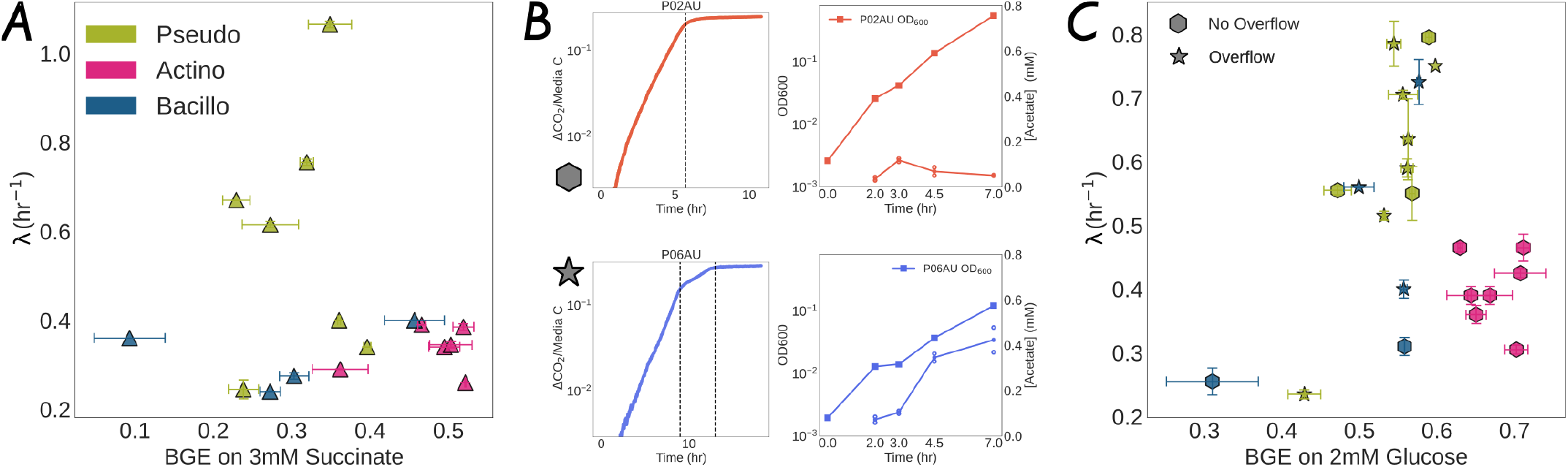
BGE is independent of growth rate. **(A)** BGE versus specific growth rate on succinate. No strain exhibits overflow on succinate. BGE and *λ* do not show a statistically significant correlation (Pearson *r* = -0.16, p-value = 0.82). Correlations between BGE and *λ* are not significant within phyla on succinate: Pseudomonadota: *r* = 0.05, p-value = 0.43, Actinomycetota: *r* = 0.23, p-value = 0.23, Bacillota *r* = 0.18, p-value = 0.32. **(B)** representative CO_2_ dynamics on glucose for two strains, illustrating single-phase respiration (no overflow, top row) and biphasic respiration (overflow, bottom row). CO_2_ dynamics for all 23 strains in this study are shown in SI Appendix, Figs. S6–S8. Right column shows OD_600_ and acetate measurements using destructive sampling during growth. These measurements were carried out during a different experiment under identical growth conditions as those that generated the CO_2_ plots. Note biphasic CO_2_ dynamics (blue, left bottom) accumulate acetate in the medium during growth. For additional acetate measurements on other strains see Fig. S11. **(C)** BGE versus specific growth rate on glucose, with strains classified by overflow status. Hexagons: no overflow (no biphasic CO_2_ dynamics). Stars: overflow. All correlations between BGE and growth rate are reported in Table 1

Before addressing the correlation between BGE and *λ* on glucose, we first report that we observed that a subset of strains growing on glucose exhibited biphasic CO_2_ dynamics, with a shift to a slower respiration phase before the final plateau (Fig. 3B, bottom row). This biphasic respiration was observed only on glucose, never on succinate. We confirmed that the cultures exhibiting the second phase were not growth limited on anything other than supplied resource carbon (See SI section 3, Fig. S10). This suggested that strains undergoing biphasic respiration on glucose were most likely undergoing overflow metabolism, followed by utilization of the accumulated acetate. Invasive measurements of acetate accumulation in the medium for a subset of strains showed that biphasic respiration corresponds to transient acetate accumulation (Fig. 3B right column, SI section 3, Fig. S11). A similar CO_2_ accumulation dynamic has been observed previously in *E. coli* (12). Thus, the endpoint BGE for overflow strains likely reflects the summed yields of biomass and CO_2_ on both substrates (see SI Section 2).

To analyze the correlation between BGE and *λ* on glucose, we partition the strains by overflow status (Fig. 3C). Overflow metabolism on glycolytic substrates like glucose produces acetate, which accumulates extracellularly and can be subsequently utilized once glucose is exhausted (12, 14). As a result, in the presence of overflow (*J*_*exc*_ *>* 0) our endpoint measurement may not quantify BGE on a single resource since both glucose and the overflow metabolite could be consumed (see SI Section 2).

No strains in the Actinomycetota phylum exhibit overflow metabolism, and there is no correlation with BGE and growth rate in this phylum. For strains exhibiting overflow metabolism, BGE and growth rate were uncorrelated for one phylum (Bacillota). However, we found that BGE and *λ* were correlated for the Pseudomonadota that exhibit overflow behavior (Table 1). We expect that this correlation is likely driven by small sample sizes (n=7).

We were able to test this hypothesis using OD estimates of biomass yield on the larger set of 62 strains examined in this study. We find that our measured BGE is strongly correlated with the measured OD yield (maximum OD/carbon supplied, Fig. S4A) on the ∼ 20 strains where BGE was quantified. This enables us to look at patterns in OD yield as a proxy for BGE across the larger library of 62 strains. In this context we cannot demarcate overflow behavior, but the Pseudomonadota do not exhibit a growth rate OD-yield correlation. And the relationship between OD yield and *λ* for 62 strains (Table S3, Fig. S3, SI section 3) is similar to the relationship between BGE and *λ* for the 23 strains presented here. We conclude that there is not a strong correlation between growth rate and BGE across strains intra-resource.

Note that when we include both resources, we find significant correlations between BGE and *λ* for Actinomycetota and Bacillota (Table 1, bottom). This inter-resource correlation occurs since both growth rate (*λ*) and BGE are higher on glucose than succinate for these two phyla (Fig. 3 A and C). In contrast, Pseudomonadota strains show higher *λ* on succinate than glucose, but show lower BGE on succinate than glucose, thus breaking the inter-resource correlation. Higher growth rates on acids for Pseudomonadota accords with their metabolic preferences for acids over sugars reported previously (17, 18).

### TCA cycle and electron transport chain gene content predicts taxonomic variation in BGE

We next tested the idea that genomic variation might be predictive of BGE across strains. Since BGE likely emerges from the balance between energy supply and demand and their coupling with CO_2_ production, we reasoned that enzymes and pathways in central metabolism might be predictive given previous studies showing that mechanistically relevant enzymes presence/absence can be predictive of bacterial phenotypes (19, 20).

We hypothesized that variation in BGE reflects genomic variation in (i) central metabolism pathways directly connected with CO_2_ production and consumption, and (ii) Electron Transport Chain (ETC) and associated redox modules that are responsible for energy production (See Table S4). To test our hypothesis, we trained a Random Forest regression model on the presence-absence of 41 KEGG-annotated genes across the aforementioned pathways on 19 sequenced strains that grew on both resources, training separately on glucose and succinate (See Fig. S18 for the genes in the feature set and their presence/absence across sequenced strains). Given the strongly conserved rank ordering of BGE on both resources across taxa, we expected to find a common set of important genes to predict BGE across resources.

The resulting predictions and gene importance scores are shown in Fig. 4. Leave-one-out cross-validation yields out-of-sample Pearson *r* ≈ 0.64 (*p* = 0.002) on glucose and Pearson *r* ≈ 0.54 (*p* = 0.04) on succinate. The two most important genes were (i) fumarate hydratase subunit beta (K01678, fumB), a TCA-cycle enzyme that does not itself release CO_2_, and (ii) the SdhD/FrdD membrane-anchor subunit (K18860, sdhD/frdD) of succinate dehydrogenase, which is simultaneously a TCA-cycle component and Complex II of the electron transport chain (succinate:quinone oxidoreductase). The KEGG ortholog does not distinguish succinate dehydrogenase from the homologous fumarate reductase; under our aerobic conditions the oxidative (succinate dehydrogenase) direction is expected. Succinate dehydrogenase does not translocate protons itself (21), unlike Complexes I, III, and IV, and so does not generate the proton-motive force directly; it reduces the quinone pool that Complexes III and IV use to pump protons, contributing to the proton-motive force indirectly.

**Fig. 4.**
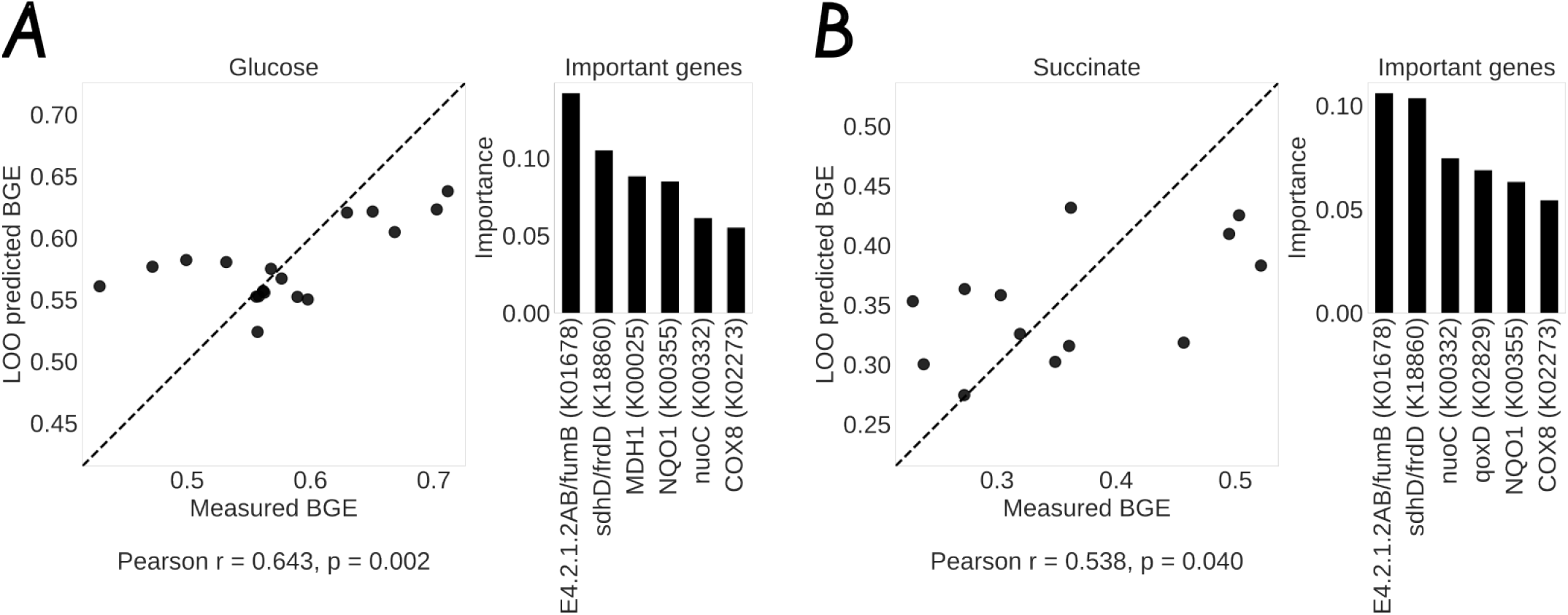
Catabolism and Electron-transport-chain genes predict BGE. **(A)** Out-of-sample fits of measured versus expected BGE for glucose after training a Random Forest model on 41 genes (or sets of genes with identical presence/absence patterns) as predictors. Genes were selected from catabolic pathways in central metabolism and the electron transport chain reflecting the importance of parameter *e*_*r*_ . Right panel shows a ranked list of the top most predictive genes. **(B)** Out-of-sample fits and ranked important genes to predict BGE on succinate. As hypothesized, we find predictive genes that are common to both resources. The top two most important genes are: (i) *fumB* (K01678), fumarate hydratase subunit beta in the TCA cycle, and (ii) a variant of *sdhD* (K18860), a subunit of succinate dehydrogenase, an enzyme shared between the TCA cycle and the ETC, functioning as Complex II in the latter.

We note that given the strong phylogenetic signature of BGE in this dataset, and its small size, a detailed mechanistic interpretation of genes detected via regression may not be warranted (20) since many genes may reliably capture phylogenetic patterns in BGE. We have attempted to limit this effect by constraining the genes used in our regression.

We also attempted to find correlations between measured BGE and genome length, as well as between BGE and genomic %GC content. We find no significant statistical correlation between BGE and genome length, while BGE correlates positively with genomic %GC content, significantly on glucose (Spearman *r*_*s*_ = 0.52, p-value = 0.026) and with a similar strength but non-significant trend on succinate (*r*_*s*_ = 0.50, p-value = 0.08). See Fig. S19. Our findings were contrary to those of Saifuddin et al.(22) who reported a negative correlation between CUE and both genome length as well as genomic %GC content.

## Discussion

BGE is hierarchically structured by substrate and taxonomy, and is independent of growth rate on strains where the endpoint measurement reflect carbon allocation. A simple stoichiometric flux model predicts the substrate ordering from canonical ATP yields. TCA and ETC gene content captures the taxonomic component of the variation, with fumarate hydratase subunit beta and succinate-dehydrogenase subunit D variants as the top two predictors on both substrates. The fate of resource carbon is thus set by catabolic machinery and biosynthetic demands, not by kinetics.

### Reconciling prior literature

Previous studies (see Table S1) have reported positive, negative, and null correlations between BGE (or CUE) and growth rate. These discrepancies fall into three categories.

Studies using indirect biomass proxies report correlations whose magnitude is comparable to the measurement uncertainty. Pold *et al*. (8) infer CUE from optical-density-based biomass estimates across a library of isolates. We find substantial variation in the ratio of OD to carbon in biomass (Fig. S4B and S4C) and the resulting uncertainties in CUE approach 100% of the mean. Similarly, Smith *et al*. (11) extrapolate biomass carbon from flow-cytometry cell-size measurements and report similarly large errors. Both studies report positive correlations whose statistical significance is difficult to assess given the measurement uncertainty.

Studies using fixed stoichiometric conversions report implausibly low BGE values. Muscarella and coworkers (10) measure BGE via leucine incorporation and oxygen evolution, converting O_2_ consumption to CO_2_ production with a fixed respiratory quotient that is known to vary with substrate (12). They report a population of strains with BGE ∼ 0.01, which purports little biomass production and high respiration and contradicts ATP balance: achieving growth rates of ∼ 0.5 h^−1^ at BGE = 0.01 would require ATP production rates an order of magnitude or more above measured maxima. This likely reflects leucine incorporation artifacts during stationary phase.

Studies reporting negative correlations rely on growth-rate estimates that exceed physiological limits. Roller *et al*. (9) report that CUE declines with growth rate across eight isolates, with *E. coli* growth rates of 2.4 h^−1^ on succinate at 25^°^C, faster than *E. coli* growth rates in rich media at higher temperatures (14).

In contrast, Andersen and von Meyenburg (12) used continuous respirometry on *E. coli* across ten carbon sources from which we find a correlation between BGE and growth rate across resources. This correlation is due to inter-resource correlations consistent with our results, since Andersen and von Meyenburg report BGE independent of growth rate on the subset of glycolytic resources, consistent with our model prediction. Our results extend this finding across three bacterial phyla and confirm that the *λ*-independence of BGE is a general feature of balanced-growth physiology, not an artifact of a single model organism.

### Implications for community carbon cycling

Ultimately, carbon cycling emerges from organic carbon utilization in a community context. Our results suggest that community-level BGE should be predictable from substrate composition and taxonomic composition, weighted by relative activities, and that growth-rate effects are secondary. ETC gene content, which is routinely recovered from metagenomic assemblies, may provide a practical genomic proxy for taxonomic BGE variation in soil and marine communities.

The large variation in BGE across taxa suggests that community composition should have substantial impacts on carbon allocation since communities dominated by, for example, Actinomycetota would be expected to exhibit higher growth efficiency. In apparent contradiction to this idea, some soil amendment studies have shown that the identity of the amended substrate has a larger effect on CO_2_ production than does origin of the soil sample (23). This discrepancy might arise from differences in the physiological state of cells in soil versus our growth experiments.

### Limitations of our study

Our study has several limitations which will be important to address in future work. First, our mathematical description of carbon allocation during aerobic growth assumes a balance between energy production via central metabolism and energy demand for growth. However, cells produce substantially more energy than can be attributed to biosynthetic demands alone. Mori *et al*. (13) recently undertook a detailed account of ATP production and utilization during balanced exponential aerobic growth in *E. coli*. They noted a major fraction of the ATP produced cannot be accounted for by the demand during growth which is assumed to be dominated by protein synthesis. This difference vanished in more energy-limited anaerobic conditions. The implication being that cells are producing far more ATP than they can consume in aerobic conditions. In the model considered here, this would have to be accounted for by inflating *σ* due to processes such as futile cycling or other energy inefficient processes. We do not understand what bounds or determines this excess energy production.

Second, our model neglected Non-Growth Associated Maintenance (NGAM) energy (*σ*_0_ [mol ATP/mol biomass C /h]). When this quantity is added to the energy demand of biomass production the result is a BGE that depends on growth rate (see SI section 1). The relevance of NGAM (*σ*_0_) to BGE drops as the ratio *λσ/σ*_0_ rises. Thus, when *σ*_0_ is a small fraction of total energy demand our result that BGE does not vary with growth rate holds. Our data do not support the existence of a universal *σ*_0_ value, which would imply slow growers have lower BGE. In fact, we observe the highest BGE values in our slow growing strains (Actinomycetota, Fig. 3A), suggesting an even smaller taxon dependent *σ*_0_. We are not aware of estimates of *λσ/σ*_0_ for non-model organisms, and even for *E. coli* estimates of *σ*_0_ can vary by a factor of *>*2.5 (Table S2).

Third, when we measured carbon left over in the supernatant (passed through a filter) we found relatively large quantities (5.7-33.3% of supplied organic carbon, Fig. S14). For a subset of strains we confirmed that this carbon was not the initially supplied resource (SI section 3 on assays, Fig. S15, S16). Therefore, we expect that this soluble organic carbon that remains in the medium has been transformed by metabolic activity and arises from the production of exopolysaccharide or excreted protein (24). However, even if we include this organic carbon in biomass the central finding of our study is unchanged (see Fig. S17).

Finally, our measurement modality provides dynamic quantification of CO_2_, but only snapshots of the other fates of carbon. The greatest impact of this limitation is that we cannot easily quantify *J*_*exc*_ (Fig. 1A). However, we show that the dynamic behavior of CO_2_ in time can reflect *J*_*exc*_ if it is taken up by cells and respired. This limitation makes it challenging for our measurement to make concrete claims about CUE as defined above.

## Conclusion

Our study substantially revises our understanding of carbon allocation during growth across diverse bacterial taxa by demonstrating that taxonomy and resource qualitatively impact the fate of carbon in a growing population. To impact our understanding of carbon cycling on a larger scale, it will be critical to extend these results to the community context, other resources such as protein or polysaccharide, and to other physiological states pertinent to wild bacterial populations.

## Materials and Methods

### Growth Phenotype Screening

A subset of 81 natural isolates (Actinomycetota: 19, Bacillota: 21, Bacteroidota: 6, Pseudomonadota: 35) from the Kuehn lab strain bank were chosen for a growth phenotype screen. The Kuehn lab strain bank consists of ≈ 400 heterotrophic strains across the 4 dominant phyla found in soil stored at − 80^°^C in glycerol. The chosen strains were revived in 0.2X Tryptic Soy Broth (TSB) for 24–48 hours at 27^°^C, in 14 mm diameter culture tubes on orbital shakers at 220 rpm. 50–100 *µ*L of the revived culture was then transferred to ≈ 5 mL of minimal M9 media (pH ≈ 7.09–7.14) with either 2 mM glucose or 3 mM succinate for 24 hours under the same temperature and shaking conditions as the previous step. M9 media for both resources had the same total carbon atom concentration i.e. 12 mM. We passaged cells from the 24 hour M9 culture to fresh M9 media for another 16-24 hour period. Finally, 500 *µ*L of the M9 culture was passaged to 4.5 mL of M9 media for 2–3 hours to bring the cells into exponential growth. 30 *µ*L of this culture was transferred to a well containing 670 *µ*L of M9 media in a 48-well lidded optical plate. The gap between the lid and plate was sealed using parafilm tape before being transferred to BMG Spectrostar Nano (platereader). Plates were cultured in the platereader at 27^°^C with 500 rpm orbital shaking in “plate mode” to obtain growth curves for strain + resource condition. OD600 measurements were taken every 5 min. The growth curves were characterized to infer maximum growth rate and max OD600 yield. During this phenotyping screen, we only considered strains that grew quantifiably on at least one resource in 40 hours. Of the 81, we successfully phenotyped 62 strains (Actinomycetota: 13, Bacillota: 18, Pseudomonadota: 31). A subset of 23 strains from the 62 successfully phenotyped strains was used for the carbon flux quantification.

### Carbon flux quantification

#### CO_2_ flux quantification in ‘Respiration Tubes’

Borosil 9900018 model glass tubes (200 mm (H) × 38 mm (W) with threaded caps) were used to culture strains for carbon quantification. Each tube, henceforth termed ‘respiration tube’, had a total volume of ≈ 182 mL. 12 mL was used for liquid culture, giving us a headspace of 170 mL. Each respiration tube was kept vertically upright on a magnetic stir plate using a custom-built rack. A 12 mm spinbar was used to aerate the culture at 500 rpm. The respiration tubes and custom-built rack were placed inside an incubator at 27^°^C. Respiration tubes were hermetically sealed using a thin layer of vacuum grease on the glass rim between the tubes and the threaded caps. The threaded caps were perforated with 2.54 mm diameter holes through which we inserted gold-plated connector pins (Male–Male). Any air gaps between the inserted pins and the perforated holes were hermetically sealed using heat-cured epoxy (EpoTek 353ND) on the topside and underside of the caps.

Sensirion SCD30 Non-Dispersive Infra-Red (NDIR) sensors were used for direct, continuous, and non-destructive measurement of gaseous CO_2_ in the headspace. 1” long F-F connector wires were used to attach SCD30 CO_2_ sensors (25.4 mm × 50.8 mm) to the underside of the caps. The sensors have an accuracy of ± 30 ppm + 3% Mean Value across a range of 400–40,000 ppm, and a fast response time ≈ 20 s. The sensors were driven by a Raspberry Pi 4B using the I2C interface. A multiplexer TCA9548A module was used to allow a single Raspberry Pi to drive up to 7 SCD30 sensors in parallel. While the TCA9548A multiplexer is an 8-channel module, we found that running all 8 channels could result in hardware failures. Using up to 4 Raspberry Pi’s, we w ere able to drive up to 28 measurements in parallel. See Fig. S20 for the circuit diagram schematic. A custom Python 3.12 script was written to collect measurements from the SCD30 sensors. Each sensor logged the CO_2_ reading in ppm, relative humidity %, and the temperature in Celsius, as well as the time of each measurement. Timestamps from all sensors were recorded as Unix epoch time to provide a common clock across our sensors and measurements. Data was stored in a CSV format for analysis. See Fig. 1B.

#### CO_2_ sensor calibration

The SCD30 NDIR CO_2_ sensors were calibrated using two methods: (i) Dilute acetic acid (CH_3_COOH) + sodium bicarbonate (NaHCO_3_) assay. The reaction of acetic acid and sodium bicarbonate releases a predictable quantity of CO_2_. We used increasing amounts of NaHCO_3_ in a hermetically sealed vessel of known volume to compare expected versus measured CO_2_ (SI section 3, Fig. S5). (ii) The SCD30 datasheet recommends a script-based calibration. The sensors were kept overnight in ambient air to equilibrate with atmospheric CO_2_ concentrations of ≈400 ppm. A function in the python script running the sensor measurements was used to override the sensor readings to report ambient CO_2_ at ≈ 400 ppm. This step was performed before each experiment to ensure consistency across sensors.

#### Inferring growth rate from CO_2_ dynamics

Carbon fluxes are assumed to be balanced during exponential growth. Thus, the CO_2_ production rate is identical to the specific growth rate. A straight line was fit to the natural log-transformed CO_2_ time series data during the exponential growth phase to obtain the growth rate (Fig. 1B). Growth rates inferred from CO_2_ dynamics are consistent with growth rates inferred from OD growth of the same strain + resource combination in the platereader under near-identical growth conditions (SI section 3, Fig. S9).

#### Converting headspace CO_2_ from ppm to moles

The ideal gas law is used to convert headspace gaseous CO_2_ readings in ppm to moles.

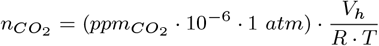

where 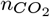 is the number of moles of gaseous 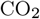 is the sensor reading, *V*_*h*_ is the headspace volume in liters (0.17 L), R is the ideal gas constant 0.0821 L·atm/(K·mol), T is the Temperature in Kelvin (300 K).

#### Accounting for Dissolved Inorganic Carbon (DIC)

The respired CO_2_ is present in the respiration tube in different forms, viz. (i) gaseous CO_2_, (ii) dissolved CO_2_, (iii) dissolved carbonate forms. To account for dissolved CO_2_, we used Henry’s law, which states that:

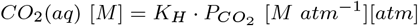

Where *K*_H_ is the Henry’s Law constant for CO_2_, 3.4 × 10^-2^ M atm^-1^, and 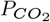 is the partial pressure of gaseous CO_2_ in the headspace in atm. The concentration of other dissolved carbonates 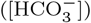 is set by the Henderson–Hasselbalch equation

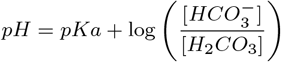

where Ka is the acid dissociation constant of H_2_CO_3_. Additionally, the TOC-L measurement described below quantifies dissolved carbonates directly. See SI section 3 and Fig. S12, S13 for additional characterization of the dissolved inorganic carbon.

#### Quantifying Carbon in the biomass and supernatant

A Shimadzu TOC-L (Total Organic Carbon Analyzer) was used to directly quantify organic and inorganic carbon in the biomass and supernatant. Carbon measurements were taken at the end of the growth phase in the respiration tubes, defined as the point when CO_2_ readings leveled off. The entire culture was removed from the respiration tube using a graduated serological pipette, and the final volume (from the original 12 mL culture) was recorded to account for water loss due to evaporation and condensation on the tube walls. The culture was transferred to a 15 mL Falcon tube. A 500 *µ*L aliquot was removed to measure OD_600_ and stored for sequencing. The remaining culture was divided into two portions. Lugol’s solution (KI + I_2_) was added to one portion (Volume A) to arrest growth without lysing bacterial cells. The second portion (Volume B) was filtered through a 0.22 *µ*m filter to separate bacterial cells (and their biomass) from the supernatant. From each portion, 4.5 mL was diluted to 18 mL in separate 24 mL glass vials compatible with the TOC-L autosampler. The TOC-L measured total carbon (TC) and total dissolved inorganic carbon (IC) via destructive high-temperature catalytic oxidation (25). Total organic carbon (TOC) was calculated as TOC = TC − IC. Biomass TOC was determined as the difference between the TOC of Volume A and Volume B, corrected for dilution.

#### Calculating Bacterial Growth Efficiency (BGE)

Bacterial growth efficiency is computed from the endpoint measurements as

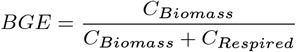

where *C*_*Biomass*_ is the utilized resource carbon allocated to bacterial biomass during the growth period, and *C*_*Respired*_ is the resource carbon respired in the same period, which consists of CO_2_ (gaseous) + CO_2_ (dissolved) + Dissolved Inorganic Carbon (DIC), the last of which is measured by the TOC-L. Under balanced exponential growth that terminates when the substrate is exhausted, this endpoint ratio is equal to the rate ratio 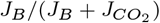 in Eq. 1 (SI Section 2). Full carbon data are provided in the supplementary dataset. See Table S5 as a guide to the full carbon dataset.

#### Information about sequenced strains

We had fully sequenced 19 of the 23 strains in this study. Strains P01AA, P03PF, P05AS, P06AU, P09PU, P11AS were sequenced for an earlier study(19). See BioProject ID PRJNA660495 for details. Strains A02AG, A03AR, A04LP, A06AU, A07AU, B01BP, B02BM, B03MU, B04BA, P02AU, P04AU, P08AV, P10PT were sequenced concurrently with this study. Approximately 5 × 10^9^ cells were preserved in DNA/RNA Shield (Zymo Research) and submitted to Plasmidsaurus (USA) for whole-genome sequencing. Genomic DNA was prepared using an amplification-free library construction protocol (ONT V14 chemistry) and sequenced on Oxford Nanopore Technology (ONT) R10.4.1 flow cells. Raw signals were base-called using the Dorado Super-Accurate (SUP) model with a default Q10 quality filter. Read quality was further refined by removing the lowest 5% of reads using Filtlong v0.2.1.

De novo assembly was executed via the Autocycler pipeline, which generates a consensus from multiple independent assemblers, including Flye v2.9.6+, Hifiasm, and Plassembler v1.8.0+. The final consensus assembly was polished using Medaka v1.8.0 and oriented to the dnaA start position using dnaapler. Structural and functional annotation was performed using Bakta v1.11.4 utilizing the v6.0 full database. Species identification was performed using Sourmash v4.6.1 against RefSeq and GenBank databases, while genome completeness and contamination were assessed via CheckM v1.2.2.

#### Building the KEGG Ortholog table

Open reading frame (ORF) prediction and functional annotation were performed on all sequenced strains using the computational tool Prodigal (version 2.6.3), for ORF-calling. Prodigal-generated output included nucleotide and corresponding amino acid sequences for each predicted gene, which served as input for downstream functional profiling. To assign genes to known functional gene families and metabolic pathways, we employed KofamScan (version 1.3.0), a profile hidden Markov model-based tool that uses KEGG Ortholog (KO) profiles to detect homology against the KEGG database. KofamScan was run with default thresholds and adaptive score cutoffs to ensure high-confidence KO assignments for each amino acid sequence. From the resulting KO annotations, we extracted KO identifiers relevant to bacterial electron transport chain, TCA cycle, and other relevant modules in core metabolism (See Table S4) and compiled them into a matrix representing presence/absence (or abundance, where applicable) of electron transport-associated KOs across all 19 strains. This KO table was used for comparative analyses of electron transport system composition among strains, yielding an initial set of approximately 240 KOs across the 19 strains. We wanted to perform a Random Forest regression using these KOs to infer the most important KOs to predict BGE across the sequenced strains in our carbon quantification dataset. However, since the size of the predictor set (number of KOs) substantially exceeded the number of observations (19 strains), the KO set was trimmed to reduce the dimensionality prior to machine learning analysis. A prevalence-based filtering approach was used to remove any KOs present in less than 2 and more than 17 strains, since we have sequenced data for 19 strains. This resulted in a much smaller KO table with 41 KOs (Fig. S18).

#### Random Forest Regression

To infer the relative importance of KOs in explaining variation in BGE, we implemented a Random Forest regression model using Python (scikit-learn 1.6.1). The truncated KO table (41 predictors) served as input features, and BGE was treated as a continuous response variable. Given the relatively small sample size ( ≈ 20 strains per resource condition), we used leave-one-out cross-validation (LOOCV) to optimize model hyperparameters. Hyperparameter tuning was conducted using the built-in model selection utility: GridSearchCV, with mean squared error (MSE) used as the performance metric. The hyperparameter grid included, the number of trees (n_estimators): [50, 100, 200], maximum tree depth (max_depth): [1, 2, 4, None], minimum samples per leaf (min_samples_leaf): [1, 2, 3, 4], minimum samples per split (min_samples_split): [2L, 3L, 4L], where L is the hyperparameter value for min_samples_leaf. The optimal parameter set was selected as the configuration minimizing cross-validated MSE. After identifying the optimal hyperparameters, we performed LOOCV prediction to generate out-of-sample predictions for each observation. These predictions were used to calculate the out-of-sample Pearson-*r* and permutation p-value. Feature importance scores were extracted from the optimized Random Forest model using the default impurity-based importance (mean decrease in impurity). KOs were ranked according to their importance scores to identify candidate ETC functions most strongly associated with BGE variation. To assess whether the observed model performance exceeded that expected under a null hypothesis of no association between KO composition and BGE, we conducted a bootstrap permutation test. Specifically, BGE values were randomly shuffled across observations while keeping the KO feature matrix fixed. For each of 1,000 iterations, the full modeling pipeline (including LOOCV and *R*^2^ calculation using the optimized hyperparameters) was repeated to generate a null distribution of out-of-sample *R*^2^ values.

## Supporting information

CO2 dynamics

Carbon quant data

## ACKNOWLEDGMENTS

S.K. acknowledges the National Institute of General Medical Sciences R01GM151538, the National Science Foundation through the Center for Living Systems (grant no. 2317138), the National Institute for Mathematics and Theory in Biology (Simons Foundation award MP-TMPS-00005320 and National Science Foundation award DMS-2235451); and the National Science Foundation CAREER award (BIO/MCB 2340416). S.K. and V.S. acknowledge funding by the Institute for Climate and Sustainable Growth at the University of Chicago. S.K. and V.S. thank Freddy Bunbury for the sequencing work on 13 of the 19 fully sequenced strains in this study.

## Notes

### Competing Interest Statement

The authors have declared no competing interest.

### Summary of Updates

Updated Fig 2A for clarity. Dataset removed while manuscript is under review.

