## Supplementary Information for "Hierarchical control of bacterial growth efficiency by substrate and taxonomy"

#### This PDF file includes:

Supporting text
Figs. S1 to S20
Tables S1 to S5
SI References

### Supporting Information Text

### Section 1: Formalism and theory of BGE

The main text defines carbon use efficiency (CUE) and bacterial growth efficiency (BGE) as rate ratios during balanced exponential growth and predicts BGE from a minimal flux model in which carbon is partitioned between biosynthesis and respiration. This section gives the mass-balance derivation of the two ratios, establishes the equivalence condition BGE = CUE, and develops the flux model.

**Mass balance and definitions.** Consider a chemoheterotroph under balanced exponential growth on a single reduced carbon source $C$, with biomass $x$, excreted inorganic carbon ($CO_2$) $C_I$, and other excreted organic compounds $C_j$. All quantities are expressed as moles of carbon atoms per unit volume. Mass conservation gives

$$\dot{C} = \dot{x} + \dot{C}_I + \sum_j \dot{C}_j. \quad [1]$$

Dividing through by biomass $x$ defines fluxes per unit biomass carbon, which we denote by $J$:

$$J_{C,\text{in}} = -\dot{C}/x, \quad [2]$$

$$J_B = \dot{x}/x = \lambda, \quad [3]$$

$$J_{CO_2} = \dot{C}_I/x, \quad [4]$$

$$J_{\text{exc}} = \sum_j \dot{C}_j/x, \quad [5]$$

where $\lambda$ is the specific growth rate. Mass balance becomes

$$J_{C,\text{in}} = J_B + J_{CO_2} + J_{\text{exc}}. \quad [6]$$

Carbon use efficiency and bacterial growth efficiency are defined as

$$\text{CUE} = \frac{J_B}{J_{C,\text{in}}}, \qquad \text{BGE} = \frac{J_B}{J_B + J_{CO_2}}. \quad [7]$$

**Equivalence condition.** From Eq. 6 and Eq. 7,

$$\text{CUE} = 1 - \frac{J_{CO_2} + J_{\text{exc}}}{J_{C,\text{in}}}, \qquad \text{BGE} = 1 - \frac{J_{CO_2}}{J_B + J_{CO_2}}. \quad [8]$$

When $J_{\text{exc}} = 0$, $J_{C,\text{in}} = J_B + J_{CO_2}$ and the two ratios are identical:

$$\text{BGE} = \text{CUE} \iff J_{\text{exc}} = 0. \quad [9]$$

When $J_{\text{exc}} > 0$, BGE exceeds CUE by a margin that depends on the magnitude of $J_{\text{exc}}$.

**Summary of prior literature and findings.** A summary of key studies and methods that report BGE, CUE, and other related quantities is shown in Table S1.

**Minimal flux model.** Under balanced growth with negligible excretion ($J_{\text{exc}} = 0$), BGE is set by two stoichiometric parameters: the energetic requirement, $\sigma$ (mol ATP / mol biomass carbon), and the net ATP yield per mole $CO_2$ respired, $e_r$ (mol ATP / mol $CO_2$). The model is adapted from (1, 2). For simplicity, we convert energy from reducing equivalents like NADH, $FADH_2$, etc. into ATP units assuming standard conversion factors.

**Energy balance** Energy produced is energy required:

$$J_E = \sigma\lambda, \quad [10]$$

where $\lambda = J_B$ is the specific growth rate, i.e. the specific carbon flux towards biomass.

**ATP yield from respiration.** The energy flux is related to the respiratory carbon flux by

$$J_E = e_r J_{CO_2}, \quad [11]$$

Combining Eqs. 10 and 11:

$$J_{CO_2} = \frac{\sigma\lambda}{e_r}. \quad [12]$$

**BGE.** Substituting into Eq. 7 with $J_{\text{exc}} = 0$:

$$\text{BGE} = \frac{J_B}{J_B + J_{CO_2}} = \frac{\lambda}{\lambda + \sigma\lambda/e_r} = \frac{1}{1 + \sigma/e_r} \quad [13]$$

BGE depends on the stoichiometric parameters $e_r$ and $\sigma$, and is independent of growth rate $\lambda$. Under $J_{\text{exc}} = 0$, BGE = CUE and the same expression describes both.

| Quantity | Definition | System | Reported range | Method | Study |
|---|---|---|---|---|---|
| | | *Pure-culture and isolate studies* | | | |
| Yield / CUE | $\frac{C_B}{C_S}$ | Pure culture (chemotrophic, aerobic + anaerobic) | 0.01 – 0.80 | Thermodynamic correlation (Gibbs energy dissipation per C-mol biomass) | Heijnen & van Dijken 1992 (6) |
| CUE | $\frac{C_B}{C_B+C_R}$ | Soil + pure culture (glucose) | 0.4 – 0.8 | Calorespirometry (heat + $CO_2$); position-specific $^{13}$C-glucose | Dijkstra et al. 2011‡ (8) |
| GE | $\frac{\Delta C_B}{\Delta C_S}$ | Pure culture (review + conceptual model; oligotroph/copiotroph) | ∼0.2 – 0.7 | Synthesis of chemostat/batch yields; rate–efficiency tradeoff framework | Roller & Schmidt 2015 (10) |
| CUE | $\frac{C_B}{C_U}$ | Bacteria (>200 species, *in silico*) | 0.22 – 0.98 (mean 0.62) | Genome-scale constraint-based metabolic modeling (FBA) | Saifuddin et al. 2019 (15) |
| BGE | $\frac{P}{P+R}$ | 20 aquatic bacterial isolates (glucose, succinate, protocatechuate) | <0.01 – 0.32 | Bacterial production ($^3$H-leucine) and respiration ($O_2$ consumption) | Muscarella, Howey & Lennon 2020 (16) |
| CUE | $\frac{G}{G+R}$ | Soil bacterial isolates (pure culture, 3 $T$, 4 substrates) | ∼0.2 – 0.7 | Optical Density (OD) measurements converted to biomass | Pold et al. 2020 (13) |
| CUE | $\frac{G}{G+R}$ | Diverse environmental isolates (pure culture) | ∼0.2 – 0.8 | Flow-cytometry biomass-C + cumulative $CO_2$ (MicroResp) | Smith, Clegg, Bell & Pawar 2021‡ (17) |
| | | *Community and environmental studies* | | | |
| BGE | $\frac{P}{P+R}$ | Natural aquatic plankton (marine + freshwater) | <0.05 – 0.6 (median ∼0.15) | Paired production ($^3$H-leucine/thymidine) and respiration ($O_2$ consumption) | del Giorgio & Cole 1998 (7) |
| CUE (SUE) | $\frac{C_G}{C_U}$ | Soil (substrate-amended, ± warming) | ∼0.2 – 0.6 | $^{13}$C-substrate partitioned into biomass vs. respired $^{13}CO_2$ across temperatures | Frey et al. 2013‡ (9) |
| CUE | $\frac{G}{G+R}$ | Soil microbial communities | ∼0.2 – 0.6 | $^{18}$O-$H_2O$ DNA labeling + respired $CO_2$ | Spohn et al. 2016(11); Geyer et al. 2019 (12) |
| CUE | $\frac{G}{G+R}$ | Model soil communities (diversity gradient) | ∼0.2 – 0.7 | $^{18}$O-$H_2O$ incorporation into DNA + $CO_2$ via IRGA | Domeignoz-Horta et al. 2020 (14) |
| CUE | $\frac{C_G}{C_G+C_R}$ | Soil (global meta-analysis) | 0.2 – 0.5 (mean 0.33) | Meta-analysis of $^{13}$C/$^{14}$C tracing and $^{18}$O-water DNA labeling | Tao et al. 2023 (18) |

**Table S1. Key studies measuring or synthesising bacterial growth efficiency (BGE), growth efficiency (GE), or microbial carbon use efficiency (CUE), grouped by pure-culture/isolate versus community/environmental systems. Notation: $C_B$ = carbon incorporated into biomass; $C_S$ = carbon in substrate consumed; $C_U$ = carbon taken up ($C_U = C_B + C_R$ at steady state); $C_G$ = carbon allocated to growth ($\equiv C_B$); $C_R$ = carbon respired as $CO_2$; $P$ = bacterial production (biomass C produced per unit time); $R$ = respiration ($CO_2$ produced, or $O_2$ consumed, per unit time); $G$ = microbial growth (biomass C produced per unit time, $\equiv P$). Rate symbols ($P$, $R$, $G$) and pool symbols ($C_x$) yield identical dimensionless ratios provided numerator and denominator are expressed in the same units (carbon mass or mol C, per unit time if rates). All the $\frac{G}{G+R}$, $\frac{P}{P+R}$, and $\frac{C_B}{C_B+C_R}$ forms are algebraically identical; the $\frac{C_B}{C_U}$ and $\frac{C_G}{C_U}$ forms are equivalent to these only when $C_U = C_B + C_R$ (i.e. no organic excretion or storage).**

**Parameters: $\sigma$, $e_r$, and the factors that affect them.** $\sigma$ has two key components, polymerization (mainly protein) and Growth-Associated Maintenance (GAM) (2) . The polymerization cost is derived from the biomass composition of cells and the fixed per-bond cost of synthesis. At a relatively fixed proteome fraction of biomass, this value is 20 mmol ATP/gDW (2), or ≈ 0.49 mol ATP / mol biomass C, which we treat as invariant across taxa and growth conditions. However, the GAM varies by taxa and growth condition, and we know neither its upper bound nor its determinants. This is a major source of uncertainty for the value of $\sigma$. FBA modeling approaches that seek to maximize the yield do so by accounting for polymerization costs and assuming a minimal GAM. This lower bound on $\sigma$ (≈ 0.49 mol ATP / mol biomass carbon) gives us an upper bound on yield, and consequently, BGE. Under these assumptions, BGE values can be unrealistically high and do not match experimental observations. The values of $\sigma$ reported in Table S2 are usually inferred from FBA models. run under different scenarios like optimizing the yield (5), or to fit experimentally measured glucose uptake rate(2), or paired with glucose uptake rate but assumed to maximize growth yield (19).

The energy ($J_E$) produced by a growing cell is not stoichiometrically locked to the $CO_2$ respired ($J_{CO_2}$); the two are only correlated, and $e_r$ is the parameter that connects them. As a baseline, we derive $e_r$ from the textbook stoichiometry of complete oxidation, i.e. the ATP produced and $CO_2$ released when a substrate is fully oxidized. Complete oxidation of glucose yields ≈32 ATP for 6 $CO_2$, giving $e_r \approx 5.3$ mol ATP/mol $CO_2$; succinate enters the TCA cycle and yields ∼16.5 ATP per turn for 4 $CO_2$, giving $e_r \approx 4.1$ mol ATP/mol $CO_2$. This is useful to anchor the value of $e_r$ but is an idealization. First, it assumes complete oxidation, whereas the $CO_2$ actually respired by a growing cell is expected to be lower since carbon is also diverted to biomass. Under overflow metabolism on glycolytic sources, TCA flux is reduced, lowering $e_r$ (≈2 by the estimates in (1)). Second, it treats the coupling between catabolic $CO_2$ efflux and energy generation as a fixed stoichiometry. In reality $e_r$ depends on the resource, on how reducing equivalents are converted to ATP and the topology of pathways that carry the flux. We discuss these below.

The conversion of reducing equivalents like NADH, NADPH, $FADH_2$, etc. to ATP is variable, not fixed. ATP is generated both by substrate-level phosphorylation and by oxidizing reducing equivalents through the electron transport chain (ETC), where the proton-motive force generated by electron transfer drives ATP synthase. Thus we can convert moles of reducing equivalents to moles of ATP depending on the P/O of the ETC components engaged. Under canonical stoichiometry 10 $H^+$ pumped per NADH across Complexes I, III, and IV, and 3 ATP per 10 $H^+$ at ATP synthase, minus transport costs, 1 NADH ≈ 2.5 ATP and 1 $FADH_2$ ≈ 1.5 ATP. Complete oxidation of one glucose then yields 4 ATP (substrate-level), 10 NADH, and 2 $FADH_2$, i.e. $4 + (10 \times 2.5) + (2 \times 1.5) = 32$ ATP for 6 $CO_2$, and thus $e_r \approx 5.3$. These factors, however, depend on which NADH dehydrogenase is active, and on the variants an organism encodes. Mori et al. (2) show that in aerobically respiring *E. coli*, the proton-pumping NDH-I gives 1 NADH ≈ 2 ATP and 1 $FADH_2$ ≈ 2.25, whereas the non-pumping NDH-II gives 1 NADH ≈ 1.25 and 1 $FADH_2$ ≈ 1.7 ATP. With these organism- and substrate-specific values, $e_r$ for *E. coli* on glucose ranges from ≈ 3.3 (NDH-II dominant) to 5.3 (canonical). The same calculation for succinate, where one succinate gives 1 ATP + 5 NADH + 2 $FADH_2$, yields $1 + (5 \times 2.5) + (2 \times 1.5) = 16.5$ ATP for 4 $CO_2$ under canonical factors, or $1 + (5 \times 1.25) + (2 \times 1.7) = 10.65$ ATP for 4 $CO_2$ otherwise, i.e., an $e_r$ range of 2.66 to 4.1 on succinate. The resulting extended range of BGE values is shown in Fig. S2.

The resource also sets $e_r$ through pathway topology. The partitioning of carbon between biomass and $CO_2$ is not a fixed property of the substrate but emerges from which central metabolic pathways carry the catabolic flux, and hence which reactions act as $CO_2$ sources, which as sinks, and which contribute to energy production. Using $^{13}$C-metabolic flux analysis, Mendonca et al. (3) showed that glycolytic substrates (e.g. glucose) are routed through upper glycolysis and the pentose phosphate pathway generating biomass precursors along with ATP and reducing equivalents, but with relatively few decarboxylation steps.
Gluconeogenic substrates (e.g. organic acids, aromatics), by contrast, enter at the TCA cycle and must run gluconeogenesis, which consumes energy, to build upper-glycolytic precursors; this route passes through additional $CO_2$-releasing TCA reactions, raising the $CO_2$ cost per unit biomass. The pathways carrying some of these fluxes also depend on the taxa and genome (5), which further complicates predicting $e_r$ for all taxa by substrate alone.

Taken together, both $\sigma$ and $e_r$ are subject to uncertainties that are unresolved by current approaches. FBA and genome-scale methods can bound the parameters, (lower bound on $\sigma$, and stoichiometric range for $e_r$). Current methods also fit these parameters or assume their values predominantly from models with few independent empirical measurements. We explore the space of possible BGE values in Fig. S2. If we assume the largest value of $\sigma = 2.04$ reported from literature (22) and use the ideal values of $e_r = 5.33$ (glucose), and 4.1 (succinate), Eq. 13 gives us BGE values of 0.72 and 0.67 for glucose and succinate respectively. However, the range of BGE values we get are ≈ 0.3 - 0.7 and ≈ 0.1 - 0.5 for the two resources, suggesting the parameter values overestimate the BGE. Smaller values of $\sigma$ overestimate BGE even more. Fig. S2 shows a plausible range of $e_r$ and $\sigma$ needed to cover the range of observed BGE. Determining when and how changes in $e_r$ and $\sigma$ drive changes in BGE or CUE is an important avenue for future work.

**Non-Growth Associated Maintenance energy and growth rate dependence.** : A classical model of bacterial growth suggests that the ATP cost per unit biomass ($\sigma$) depends on the growth rate and includes a growth-rate independent contribution to cellular maintenance, termed Non-Growth-Associated Maintenance (NGAM) (2, 4). We ignore this term when writing eqn(10), and subsequently we predict that BGE is independent of $\lambda$. Adding the maintenance energy component modifies eqn(10) to:

$$J_E = \sigma_0 + \sigma\lambda \qquad [14]$$

where $\sigma_0$ is the contribution for maintenance (NGAM), measured in moles ATP per moles unit biomass carbon per unit time. Going through the same balance argument in our minimal flux model, we derive a new expression for BGE:

$$\begin{aligned} \text{BGE} &= \frac{J_B}{J_B + J_{CO_2}} = \frac{\lambda}{\lambda + J_{CO_2}} \\ \text{BGE} &= \frac{e_r}{e_r + \sigma + \sigma_0/\lambda} \\ \text{BGE} &= \frac{1}{1 + (\sigma + \sigma_0/\lambda)/e_r} \qquad [15] \end{aligned}$$

In eqn. 15, BGE depends on the ratio of $g(\lambda) = \sigma + \sigma_0/\lambda$ and $e_r$. This form suggests a clean separation between the magnitude of the BGE response to the magnitude of changes in $g(\lambda)$ and $e_r$ on BGE, with the relative sensitivity to both parameters is given by:

$$\frac{\partial \ln \mathrm{BGE}}{\partial \ln e_r} = \frac{g}{e_r + g} = 1 - \mathrm{BGE}$$

$$\frac{\partial \ln \mathrm{BGE}}{\partial \ln \lambda} = \frac{\sigma_0/\lambda}{e_r + g} = r(\lambda)(1 - \mathrm{BGE})$$

where:

$$r(\lambda;\, \sigma, \sigma_0) \equiv \frac{\sigma_0/\lambda}{\sigma + \sigma_0/\lambda} = \frac{1}{1 + \sigma\lambda/\sigma_0} \quad [16]$$

Both log-sensitivities factorize into a common term $(1 - \mathrm{BGE})$, which sets the overall responsiveness of BGE, and a parameter-specific weight (1 for $e_r$, and $r(\lambda)$ for $\lambda$).

Because $(1 - \mathrm{BGE})$ is shared, it cancels when we compare the two responses. The growth-rate sensitivity relative to the BGE sensitivity is then simply

$$\frac{\partial \ln \mathrm{BGE}/\partial \ln \lambda}{\partial \ln \mathrm{BGE}/\partial \ln e_r} = r(\lambda) = \frac{1}{1 + \sigma\lambda/\sigma_0}. \quad [17]$$

Thus the maintenance term influences BGE only through $r(\lambda)$. When $\sigma_0 \ll \sigma\lambda$, we have $r(\lambda) \to 0$, so the $\lambda$-sensitivity is negligible relative to the $e_r$-sensitivity, and BGE becomes effectively independent of growth rate recovering Eqn. 13.

Table S2 reports values of $\sigma_0$, $\sigma$, and $\lambda$ predominantly for *E.coli* strains. We do not report such measurements of $\sigma$ or $\sigma_0$ in our study but we speculate that $\sigma_0$ could be strain dependent. Eqn. 15 suggests that as $\sigma_0/\lambda$ increases for a given $\sigma$, BGE should decrease, i.e if $\sigma_0$ is a taxon independent quantity, slow growth should reduce BGE. However, the highest BGE values in our study correspond to slower growing strains on both resources, suggesting that the NGAM $\sigma_0$ could be taxon dependent and small for slower growing taxa, thereby once again reducing $\lambda$ dependence according to our original model.

| Organism | $\sigma_0$ (mol ATP / mol C / h) | $\sigma$ (mol ATP / mol C) | $\lambda$ (1/h) | Reference |
|---|---|---|---|---|
| *E. coli* K-12 NCM3722 | 0.49 | 1.2 | 0.8 | Mori & Hwa 2023(2) |
| *E. coli* W3110 | 0.185 | 0.32 | 0–0.65 | Varma & Palsson 1994(19) |
| *E. coli* K-12 MG1655 | 0.206 | 1.471 | – | Feist et al. 2007(20) |
| *E. coli* DH5$\alpha$ | 0.187 | – | – | Selvarasu et al. 2009(21) |
| *P. putida* KT2440 | 0.095 | 2.04 | 0–0.59 | van Duuren et al. 2013(22) |
| *P. putida* KT2440 (*i*JN1462) | 0.023 | – | – | Nogales et al. 2020(23) |
| *M. maripaludis*$^{\dagger}$ | 0.188 | 0.65 | – | Richards et al. 2016(24) |

**Table S2. Maintenance energy ($\sigma_0$) and growth-associated energy ($\sigma$) for bacteria, converted to units of mol ATP per mol biomass carbon (per hour for $\sigma_0$), with reported growth rates $\lambda$. Conversion assumes biomass is 50% carbon by dry weight (1 g DW = 1/24 mol C). Studies in this table use FBA models to set $\sigma$. $^{\dagger}$Anaerobic; shown for comparison.**

### Section 2: Inferring BGE from measurements

Our experiment measures $CO_2$ in real time, but for biomass and spent media carbon, it measures endpoint quantities, not rates. This section shows that under balanced growth the endpoint BGE ratio equals the rate-defined BGE of Eq. 7, tabulates what combinations of measurements recover CUE and BGE, and addresses the complication of overflow metabolism.

**Rate–endpoint equivalence.** We measure $C_{\mathrm{biomass}}(T) = x(T) - x(0)$ and $C_{CO_2}(T) = C_I(T) - C_I(0)$, where $T$ is a time well past the cessation of growth. The endpoint BGE is

$$\mathrm{BGE}_{\mathrm{endpoint}} = \frac{x(T) - x(0)}{[x(T) - x(0)] + [C_I(T) - C_I(0)]}. \quad [18]$$

Under balanced exponential growth, $\dot{x} = \lambda x$ and $\dot{C}_I = J_{CO_2} x$ with $J_{CO_2}$ constant, so $x(t) - x(0) \approx x(0)(e^{\lambda t} - 1)$ and $C_I(t) - C_I(0) \approx x(0)(J_{CO_2}/\lambda)(e^{\lambda t} - 1)$. Substituting into Eq. 18 and canceling the common factor $x(0)(e^{\lambda t} - 1)$,

$$\mathrm{BGE}_{\mathrm{endpoint}} = \frac{1}{1 + J_{CO_2}/\lambda} = \frac{J_B}{J_B + J_{CO_2}} = \mathrm{BGE}, \quad [19]$$

using $J_B = \lambda$. The endpoint ratio equals the rate ratio exactly under balanced growth, and differs from it by terms of order $x(0)/x(T) = e^{-\lambda T}$ when the initial biomass is not negligible. This approximation is excellent for our experiments ($e^{\lambda T} \sim 10^2$

or larger). The derivation assumes growth ceases when $C \to 0$, which is approximately correct; residual respiration during stationary phase introduces small, strain-dependent corrections.

The same argument does not apply to CUE endpoint measurements when $J_{\text{exc}} \neq 0$, because excreted compounds may be re-consumed after primary substrate exhaustion. The endpoint ratio $[x(T) - x(0)]/[x(T) - x(0) + C_I(T) - C_I(0) + \sum_j C_j(T) - C_j(0)]$ tracks CUE only if the excreted compounds remain in the medium; otherwise it tracks something between BGE and CUE (see overflow subsection below).

Our experiment measures $x(T)$, $C_I(t)$, and the full carbon budget at the endpoint. This gives BGE directly via Eq. 18; CUE is recoverable via the full mass balance.

**Endpoint BGE under overflow metabolism.** Some of the strains in this study undergo overflow metabolism on glycolytic substrates. Overflow metabolism produces acetate that accumulates extracellularly and can be subsequently re-consumed. The reconsumption process is accompanied by a second, distinctly slower, respiration phase. If the endpoint measurement is taken well after the second respiration phase is complete, the excreted acetate has been re-metabolized to biomass and $CO_2$. At the endpoint, $C_j(T) \approx 0$, and the endpoint measurement satisfies

$$C_{\text{biomass}}(T) + C_{CO_2}(T) \approx -[C(T) - C(0)] = C_0, \qquad [20]$$

since all carbon taken up has been converted to either biomass or $CO_2$. Under this condition, endpoint BGE approximately equals CUE even for overflow strains.

The effective ATP yield for overflow strains is a weighted average of the yield on glucose catabolism and the yield on acetate catabolism, with weights given by the fraction of carbon flux routed through each pathway. If the overflow fraction is a strain-level property independent of $\lambda$, BGE remains $\lambda$-independent (consistent with Fig. 3C of the main text). If instead the overflow fraction scales with $\lambda$, a weak negative correlation between BGE and $\lambda$ could appear on overflow strains; we do not observe this.

### Section 3: Additional experimental details

**Minimal defined media composition.** All strains were grown in M9 defined media background. We prepared a stock solution of 5X M9 salts that was diluted to 1X for the final media. Composition of M9 5X salts (from Difco): Disodium Phosphate (anhydrous) 33.9 g/L, Monopotassium Phosphate 15.0 g/L, Sodium Chloride 2.5 g/L, Ammonium Chloride 5.0 g/L. We added either glucose or succinate to each 1X M9 background, ensuring that the total resource carbon concentration in either media was 12 mM C atoms. For glucose + M9, we added 2 mM glucose, and for succinate + M9 we added 3 mM succinate. The pH of the media was titrated to 7.09-7.14 for both carbon resources. We measured the media carbon concentration for each batch of media carbon quantification experiments using the Shimadzu TOC-L.

**Platereader phenotyping results.** We successfully characterized 62 strains from our isolate strain bank: Pseudomonadota phylum: 31, Actinomycetota: 13, Bacillota: 18. Strains were analyzed using python scripts to smooth noisy data, infer the growth rate using a spline fit to calculate the derivative over a moving window in the exponential phase. The OD yield was calculated by taking the difference between the minimum and maximum values of path-length corrected optical density absorbance measurements at 600 nm ($OD_{600}$, which we use interchangeably with OD in this manuscript). The ranked OD yields across the strains and resources are shown in Fig. S3A. Correlations between the OD yield and growth rate are shown in Table S3 and scatter plots are shown in Fig. S3B. (A strain was considered successfully characterized if it showed a clear exponential growth phase and at least a 3-fold change in OD over time.)

The pattern of correlation between BGE and growth rate we report in the main text after analyzing our carbon quantification is consistent with our OD measurements across this larger dataset (Table S3). We observed no significant correlation between OD yield, which is analogous to BGE, and growth rate for Pseudomonadota and Bacillota phyla on both resources. We see a weak positive correlation between growth rate and yield for Actinomycetota on glucose. We observe no significant correlation across all strains for either resource, and for the whole dataset, lumping together strains and resources. We also report no significant correlation for Pseudomonadota and Bacillota strains across both resources. We do see a weak positive correlation for Actinomycetota strains.

We do not identify strains undergoing overflow metabolism due to the experimental limitations of our phenotyping approach (growth plates were not destructively sampled for acetate assays and it is difficult to identify overflow stages through OD dynamics alone).

**OD yield versus BGE and biomass carbon.** We measured the OD yield for each strain-resource pair across our carbon quantification dataset. We plotted the OD yield against both the BGE and the measured biomass carbon (Fig. S4A and B). The 2 pairs of quantities are very strongly positively correlated; we observe a resource-dependence effect. We compute the biomass Carbon concentration per OD yield (C/L/OD) across strain and resource pairs and report a mean value of 180.29 mg C/L/OD for biomass carbon, and found a non-negligible coefficient of variation of the biomass carbon/OD unit across our measurements (32%) Fig. S4C.

| | | Pearson $r$ | $p$-value |
|---|---|---|---|
| *By carbon source and phylum* | | | |
| Succinate | Pseudomonadota | 0.058 | 0.404 |
| | Actinomycetota | 0.252 | 0.210 |
| | Bacillota | − 0.170 | 0.736 |
| Glucose | Pseudomonadota | 0.020 | 0.468 |
| | Actinomycetota | 0.510 | 0.052 |
| | Bacillota | 0.090 | 0.373 |
| *By resource condition (all phyla pooled)* | | | |
| Succinate | | − 0.152 | 0.854 |
| Glucose | | − 0.136 | 0.851 |
| Overall | | − 0.099 | 0.851 |
| *By phyla (inter-resource)* | | | |
| Pseudomonadota | | −0.028 | 0.579 |
| Actinomycetota | | 0.510 | 0.005 |
| Bacillota | | 0.131 | 0.229 |

**Table S3. Pearson correlations between OD yield and $\lambda$. Top: by phylum and carbon source. Middle: By resource, pooled across phyla. Bottom: By phyla, pooled across resources. p-values are computed from a one-sided permutation test ($10^5$ shuffles) against the null of no positive correlation between OD yield and $\lambda$.**

**$CO_2$ sensor calibration.** We used two methods to calibrate the Sensirion SCD30 NDIR $CO_2$ sensors. The first method was a chemical calibration based on the reaction of sodium bicarbonate with dilute acetic acid that releases a stoichiometrically defined quantity of $CO_2$. The chemical reaction is $\mathbf{NaHCO_3 + CH_3COOH \rightarrow CH_3COONa + H_2O + CO_2}$. We used this reaction to build a calibration curve for a randomly chosen sensor. We prepared a hermetically sealed vessel (volume 260 mL) with a fixed but small quantity (10 mL) of 10–20% dilute acetic acid. We ensured that the reaction was limited by the amount of $NaHCO_3$, thereby ensuring that the moles of $CO_2$ on the product side were equal to the moles of $NaHCO_3$ added as reactants. We added increasing amounts of $NaHCO_3$ (0.05, 0.1, 0.2, 0.3 millimoles) before quickly sealing the vessel and gently agitating the solution to speed up the release of $CO_2$. The agitation and low pH of acetic acid ensure that almost all of the produced $CO_2$ is in the gaseous phase in the vessel headspace. The $CO_2$ measured by the sensors is plotted against the expected $CO_2$ in Fig. S5. The equation to convert moles of $CO_2$ to ppm readings is as follows:

$$ppm_{CO_2} = \left( n_{CO_2} \frac{\cdot R \cdot T}{V_h} \cdot 10^6 \right) + \text{Background}$$

where $n_{CO_2}$ is the number of moles of gaseous $CO_2$, $ppm_{CO_2}$ is the sensor reading, $V_h$ is the headspace volume in liters (0.25 L), R is the ideal gas constant 0.0821 L·atm/(K·mol), T is the Temperature in Kelvin (300K), and 'Background' is the atmospheric background reading of ambient $CO_2$ (400–500 ppm).

Second, we followed the manufacturer-recommended field calibration procedure described in the Sensirion specification sheet: 'Field Calibration for SCD30'. In this approach, sensors were exposed to ambient air for a minimum of 8h, after which a single command triggered an internal calibration to align the sensor readings with the ambient $CO_2$ concentration (assumed to be 400 ppm). This calibration step was performed prior to each experiment to minimize signal drift and ensure measurement consistency across the whole dataset.

**Correlation between inferred growth rates from platereader OD and $CO_2$ measurements.** The $CO_2$ dynamics for all strains in this study are shown in Fig. S6 - S8. Growth rates were determined using two independent measurement approaches: OD over time during our phenotyping step and $CO_2$ dynamics during carbon quantification experiments. Because the two methods rely on distinct measurement principles and were conducted using different experimental apparatus, differences in absolute growth rate estimates were expected. However, we found a correlation between the two measured growth rates of approximately 0.7 (Fig. S9).

**Yield on glucose is limited by carbon.** On a subset of strains growing on glucose, we observed a second, slower respiration phase (biphasic respiration). See Figs. S6 - S8. This suggested either overflow metabolism leading to acetate production followed by utilization or resource-limitation to growth other than carbon that caused the extant biomass to respire the remaining carbon resource without growth. To verify that biomass yield on glucose was only carbon-limited under the experimental conditions, we cultured a subset of strains in media with increasing glucose concentrations in M9 minimal medium. Six strains were selected in total across the three dominant phyla, including four strains that exhibited a secondary respiration phase in $CO_2$ production and two that did not. In the experiments presented in the main text, cultures were grown in 2mM glucose supplemented M9 medium. For validation of our carbon-limited hypothesis, strains were pre-cultured under these same conditions and subsequently inoculated into media containing either 2 mM or 4 mM glucose in 48-well lidded optical Nunc plates. Cultures were incubated at 27°C with continuous shaking at 500 rpm. OD measurements were taken every 5 minutes. Growth under these conditions indicated that biomass accumulation at 2 mM glucose was limited by carbon availability, resulting in higher yields following identical dynamics at higher glucose concentrations. The growth curves for these experiments are shown in Fig. S10.

**Acetate Assay and Overflow Metabolism.** To test the biphasic respiration due to overflow metabolism hypothesis from the main text, we quantified the acetate production during growth on glucose. A subset of six strains was selected for analysis, four strains exhibiting biphasic respiration (*P01AA, P06AU, B01BP, B03MU*) and two strains that do not exhibit biphasic respiration (*P02AU, A02AG*). We cultured the strains in near-identical conditions as the carbon quantification, i.e. in respiration tubes with spin-bars, however, without a $CO_2$ sensor. At defined time points along the growth curve, the OD for each culture was measured, and 1 mL of culture was harvested and frozen at -20°C for further processing. Samples were subsequently thawed and then centrifuged to pellet cells, and the supernatant was collected for acetate quantification. Acetate concentrations were measured using a BioAssay Systems EnzyChrom acetate assay kit, a colorimetric assay by measuring absorbance at 570 nm. We extended the sensitivity of the assay by increasing the incubation time from 30 min to 150 min at 25°C and using 96-well half-area plates to increase the optical path length for the assay's reaction volume. A standard curve was generated for each assay to enable accurate quantification of acetate levels. We ran two assays using the kit. In the first assay, experimental samples were run alongside a seven-point calibration series (1.0 - 0.016 mM, twofold serial dilutions); however, the resulting calibration curve was non-monotonic at several points, rendering it unsuitable for quantification. A second assay was therefore performed using a fresh kit, running only the calibration standards under identical incubation and colorimetric development conditions. To account for background absorbance differences between the two kit batches, an inter-assay offset was calculated as the difference between the minimum absorbance values observed in each assay (offset = min(Assay 1) - min(Assay 2)). This offset was added to all absorbance values from the second assay prior to curve fitting, effectively aligning the baseline of the new calibration curve to that of the first assay. The offset-corrected calibration curve from the second assay was then used to calculate acetate concentrations for the experimental samples measured in the first assay (Fig. S11A and B).

We observed that the strains exhibiting biphasic respiration produced extracellular acetate, while strains without biphasic respiration did not, consistent with our overflow metabolism hypothesis.

**Calculated and measured dissolved inorganic carbon.** The respired $CO_2$ exists across different pools in our culture device. If $\Delta n_{CO_2}$ are the total moles of respired inorganic carbon, the partitioning is as below:

$$\Delta n_{CO_2} = n_{CO_2}(gas) + n_{CO_2}(aq) + n_{Carbonates}$$

where $n_{CO_2}(gas)$ : moles of $CO_2$ gas in headspace, $n_{CO_2}(aq)$: moles of dissolved $CO_2$ in aqueous culture, $n_{Carbonates}$: moles of other dissolved carbonates ($HCO_3^{-1}$, $CO_3^{-2}$)

$$n_{CO_2}(gas) = ppm_{CO_2} \cdot 10^{-6} \cdot 1atm \cdot \frac{V_h}{R \cdot T}$$

where $ppm_{CO_2}$ is the background subtracted $CO_2$ sensor reading. From Henry's Law:

$$n_{CO_2}(aq) = ppm_{CO_2} \cdot 10^{-6} \cdot 1atm \cdot K_h \cdot V_l$$

where $K_h$ is the Henry's law constant for $CO_2$ (0.034 mol/L atm) and $V_l$ is the volume of the culture in liters (0.012 L) Using the Henderson–Hasselbalch equation, we can calculate the moles of dissolved carbonates at the pH of the culture.

$$n_{Carbonates} = n_{CO_2}(aq) \cdot 10^{pH-pKa1} \quad [21]$$

where, pH is the pH of the culture and pKa1 is the first dissociation constant of $H_2CO_3$ (6.36). Thus, the total moles of respired inorganic carbon are given by:

$$\Delta n_{CO_2} = ppm_{CO_2} \cdot 10^{-6} \cdot 1atm(\frac{V_h}{R \cdot T} + K_h \cdot V_l(1 + 10^{pH-pKa1})) \quad [22]$$

(The background atmospheric $CO_2$ also gets partitioned across these pools, but the effect is negligibly small and is not due to bacterial respiration.) Equation 22 allows us to calculate the amount of inorganic carbon in dissolved carbonate form (DIC) using the $CO_2$ concentration in the headspace measured by the NDIR sensors. However, we also measure the concentration of dissolved carbonates in the culture directly using a Shimadzu TOC-L instrument. To make this measurement, the TOC-L uses 1M HCl to substantially lower the pH of the sample, shifting the carbonate equilibrium toward $CO_2$(aq). The instrument then sparges the sample with high-purity, $CO_2$-free gas to obtain a gas sample for $CO_2$ concentration measurement. We compare the measured and predicted values of dissolved inorganic carbon in Fig. S12.

A subset of points in Fig. S12 falls along the dotted "Calculated = Measured" line, indicating good agreement between the directly measured DIC concentration and the value calculated from headspace $CO_2$ via Equation 22. A second subset, however, shows systematic disagreement, with the measured DIC concentration consistently lower than predicted. This discrepancy exhibits a clear phylogenetic signal, with disagreement concentrated especially among strains from the Actinomycetota phylum, rather than distributed randomly across strains. We considered four possible sources of this discrepancy: (i) error in the TOC-L measured DIC concentration, (ii) error in the pH used in the carbonate equilibrium calculation, (iii) error in the NDIR-derived $CO_2$ readings used as an input to the calculation, or (iv) batch effects due to different durations of experiments. To evaluate (ii), we asked what pH would be required to reconcile the measured and calculated DIC values. This back-calculation yields a required pH of ≈6.4, substantially lower than our measured pH of ≈6.94. A pH this low is also inconsistent with the experimental setup, as the growth medium is buffered with a starting pH of 7.09-7.14, making a shift to 6.4 unlikely. To evaluate (iii), we used the measured DIC concentration to back-calculate the implied $CO_2$ concentration in the headspace. The resulting values are substantially smaller than the NDIR-measured $CO_2$ concentrations, and the resulting carbon mass balance fails (e.g. we cannot account for the added carbon in the medium). Having ruled out pH error and finding that $CO_2$ sensor error would itself produce an inconsistent mass balance, we conclude that the discrepancy most likely reflects a genuine loss of dissolved inorganic carbon for this subset of strains, potentially through a biological mechanism specific to particular taxa, consistent with the observed phylogenetic signal. One possible mechanism is biomass-associated inorganic carbon that stays locked in the cells and is not affected by the pH-lowering measurement method. We do not further investigate the mechanistic basis of this loss here. Finally, we assessed the impact of this discrepancy on our principal results by recalculating BGE using the predicted DIC concentration based on the headspace $CO_2$ measurements. This substitution has a negligible effect on the resulting BGE estimates and does not alter the relative ranking of BGE across strains, indicating that the discrepancy does not affect our main conclusions; see Fig.S13A for the calculated DIC values and Fig.S13B for a comparison of the BGE values.

**Resource leftover in the spent media.** Fig. S14 shows the relative proportions of the different carbon pools at the end of the growth phase as measured by our methods. The final extracellular organic carbon in the spent media (or Supernatant TOC) when normalized to the initially provided resource carbon spans a range from 5.7-33.3% of the initially provided carbon. We expect this carbon pool to consist of some original resource carbon, some cellular exudate, and a fraction of dead cell debris. To determine what fraction of the original resource is still left in the spent media, we ran metabolic assays on the spent media. We picked 8 strains across the 3 phyla and across a range of supernatant TOC concentrations. We used a Cell BioLabs STA-680 metabolic assay kit for quantifying glucose, and an EnzyChrom ESNT-100 assay for quantifying succinate (colorimetric method). We find that there is a negligibly small fraction of the original resource left in the spent media. Results for both resources are shown in Fig. S15 (Glucose) and Fig S16 (Succinate). Studies have shown that organic exudates during bacterial growth can form a significant fraction (15-20%) of the original resource carbon (26). Even if we treat this exudate fraction as a part of the biomass, it does not change our central findings. To do so, we add the spent media organic component to biomass and redo the BGE calculation. The results of this analysis are shown in Fig. S17.

**Predicting BGE from genomic information.** Fig. S18 shows the gene presence-absence table we built to find genomic features that can predict BGE using a Random Forest model. We focused on genes from modules and pathways essential for core metabolism and energy production via the electron transport chain, especially in their connection with being sources or sinks of $CO_2$. See Table S4. We also try to find a correlation between BGE and genome length or genomic %GC content. We find a moderate positive correlation between BGE and %GC content, but we find no correlation between BGE and genome length, contrary to the findings of (15).

**Circuit diagram schematic for $CO_2$ measurement hardware.** The wiring diagram (circuit diagram) for connecting a Raspberry Pi 4B to multiple Sensirion SCD30 $CO_2$ sensors is shown in Fig. S20.

**Carbon Quantification Table Description.** All the measured carbon components are available in the accompanying dataset. File: SinhaKuehn_CQuantData.csv. Descriptions of column fields for the attached data are given in TableS5. All the $CO_2$ dynamics data for each strain are attached as SinhaKuehn_CO2Dynamics.zip. Each filename in the zip file has a strain name, a resource identifier (G: glucose, S: succinate), and a replicate number (1 or 2).

| # | Module / pathway | Role |
|---|---|---|
| 1 | TCA cycle | Source |
| 2 | Pyruvate dehydrogenase (PDH) | Source |
| 3 | Glyoxylate shunt | Sink |
| 4 | Anaplerosis | Sink |
| 5 | Gluconeogenesis ($CO_2$ release on gluconeogenic substrates) | Source |
| 6 | Oxidative pentose phosphate pathway (ox-PPP) | Source |
| 7 | Electron transport chain (ETC) | Indirect |
| 8 | NDH-I and NDH-II (NADH dehydrogenases) | Indirect |
| 9 | Terminal oxidases | Indirect |
| 10 | Overflow metabolism | Source |
| 11 | Fatty acid biosynthesis | Sink |

**Table S4. Metabolic pathways relevant to bacterial $CO_2$ production and consumption and those that play a role in ATP (energy) production. Role: *source* = net $CO_2$ producer;*sink* = net $CO_2$ consumer; *indirect* = affects $CO_2$ flux via redox balance rather than direct decarboxylation, partly coupled with ATP production.**

| # | Column | Description |
|---|---|---|
| 1 | Strain Code | Strain identifier/code used in the manuscript |
| 2 | Strain | Internal strain name notation for code attached to manuscript |
| 3 | Phyla | Taxonomic phylum for the strain |
| 4 | Genus | Genus and species (if available) information on strain |
| 5 | Resource | Carbon source used in the experiment (either Glucose or Succinate) |
| 6 | Condition | Experimental condition or treatment label (<strain>_<resource letter>_<replicate number>) |
| 7 | Media C | Carbon concentration provided in defined media. Measured using TOC-L (mg/L) |
| 8 | Cul TC | Total carbon in liquid culture at the end of growth measured using TOC-L (mg/L) |
| 9 | Cul IC | Inorganic carbon in liquid culture at the end of growth measured using TOC-L (mg/L) |
| 10 | Cul TOC | Total organic carbon in the culture (Cul TC - Cul IC) (mg/L) |
| 11 | Sup TC | Total carbon in 0.22$\mu$m filtered spent media (mg/L) |
| 12 | Sup TOC | Total organic carbon in 0.22$\mu$m filtered spent media (Sup TC - Cul IC) (mg/L) |
| 13 | Bio | Biomass carbon at the end of growth (Cul TC - Sup TC) (mg/L) |
| 14 | CO2mgL | Headspace $CO_2$ measured by NDIR sensors converted to equivalent dissolved C in culture volume (mg/L) |
| 15 | BGE | BGE computed from carbon measurements |
| 16 | CO2 growth rate | Growth rate inferred from NDIR sensor measured $CO_2$ dynamics (1/h) |
| 17 | OD Growth Rate | Growth rate inferred from Optical Density measurements in platereader (separate experiment for the same strain and resource) (1/h) |
| 18 | OD600 | $OD_{600}$ yield of carbon quantification culture |
| 19 | Overflow rate | Inferred rate of respiration on overflow products from NDIR $CO_2$ sensors (1/h) |
| 20 | pH | Measured culture pH at the end of growth in carbon quantification experiment |

**Table S5. Description of columns with attached data file containing carbon quantification**

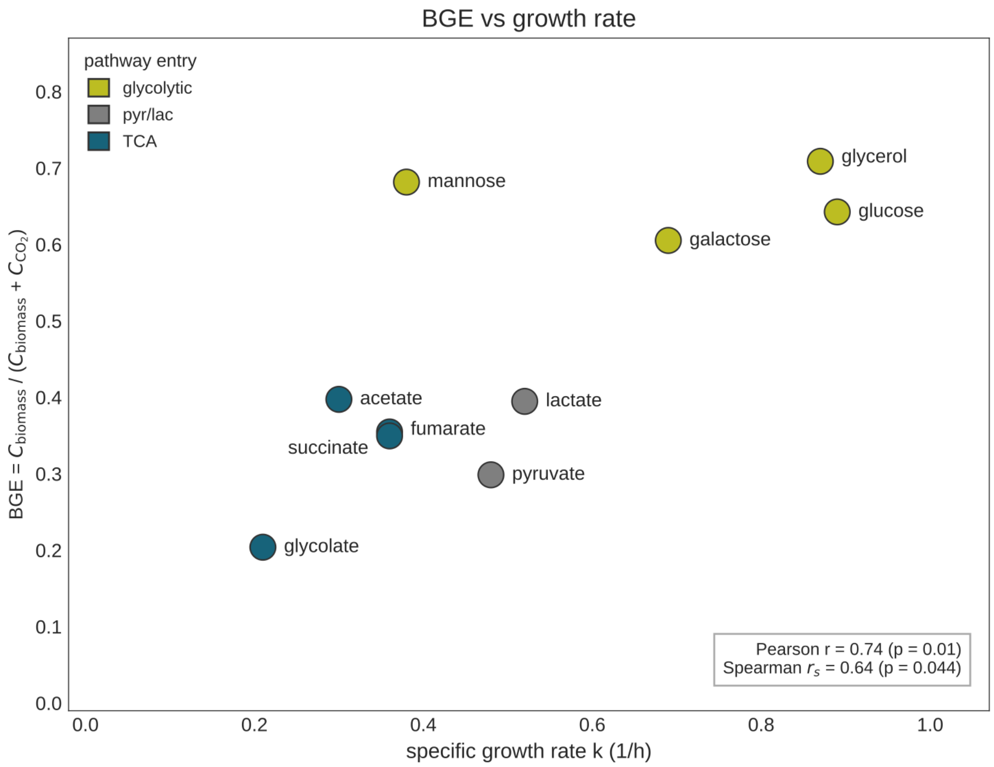


**Fig. S1.** Plotting BGE versus growth rate for *E.coli* from data in Andersen *et al.*(25). The study does not report BGE directly, so we infer it from the reported data.

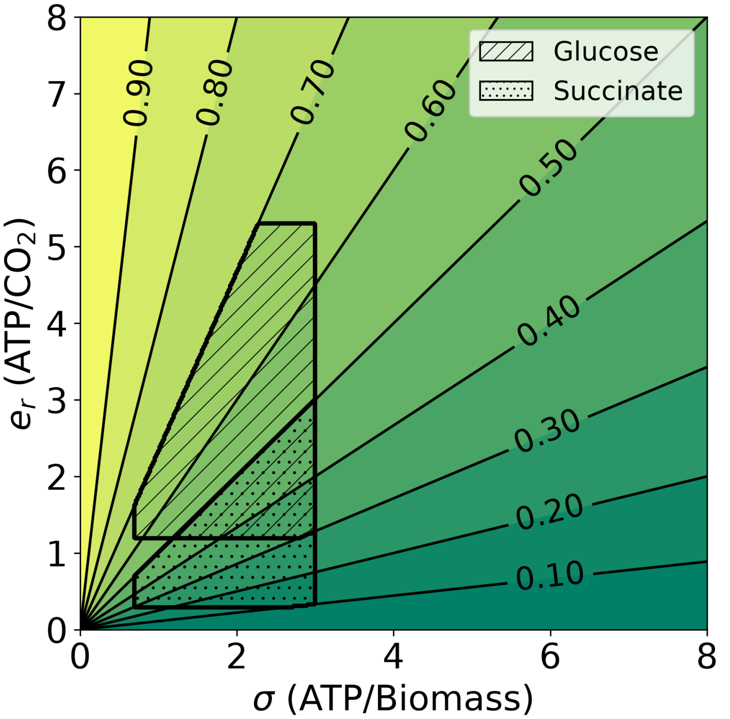


**Fig. S2.** Predicted BGE as a function of ATP yield $e_r$ and energetic ATP demand $\sigma$ (Eq. 13). The contour lines indicate ridges of equal BGE. Shaded regions indicate the span of $e_r$ and $\sigma$ to cover the BGE values reported in this study, while being arbitrarily anchored to values of $\sigma$ and $e_r$ that seem plausible based on simple assumptions outlined in the SI.

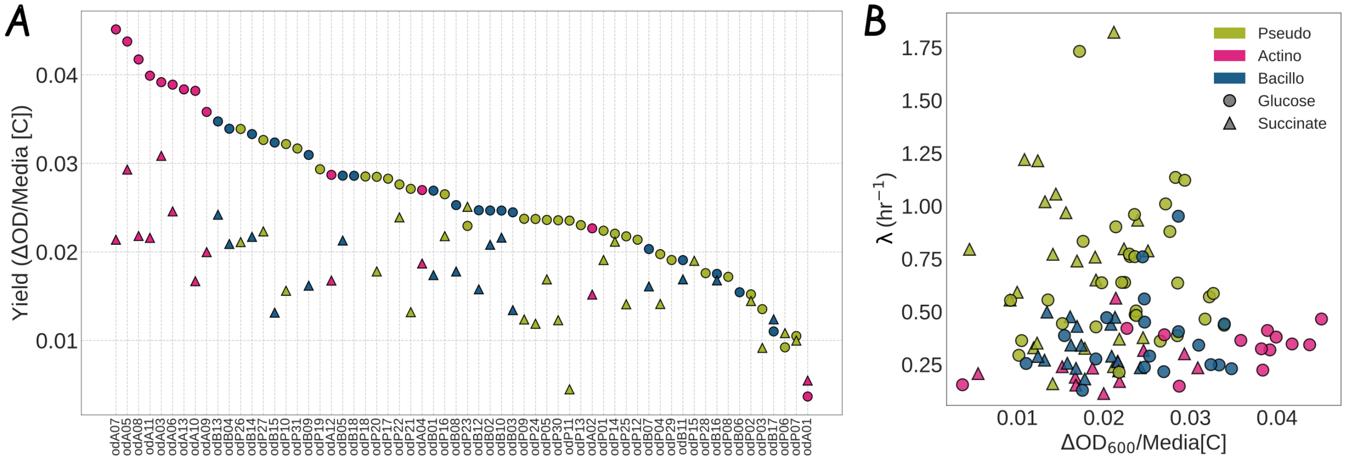


**Fig. S3.** **(A)**:Ranked OD yield (Y-axis) across 62 strains (X-axis). In 58 cases, the OD yield on glucose (circles) is higher than on succinate (triangle). In the remaining 4 cases, the difference in the yields is small and could be obscured by the noise in the OD data. Ranked correlation metric between the yields on both resources Spearman $r_s$:0.57, p-value: $\approx 10^{-5}$. **(B)**: OD yield versus growth rate for 62 strains (Pseudomonadota: 31, Actinomycetota: 13, Bacillota: 18) across two resources: glucose and succinate. Yield = $\Delta$OD/Media [C] in mM. Circular markers: glucose, triangular markers: succinate. Markers are colored by the phyla the strains belong to. Correlations in table S3. Plotted mean values of OD yield and growth rate for 2 biological replicates.

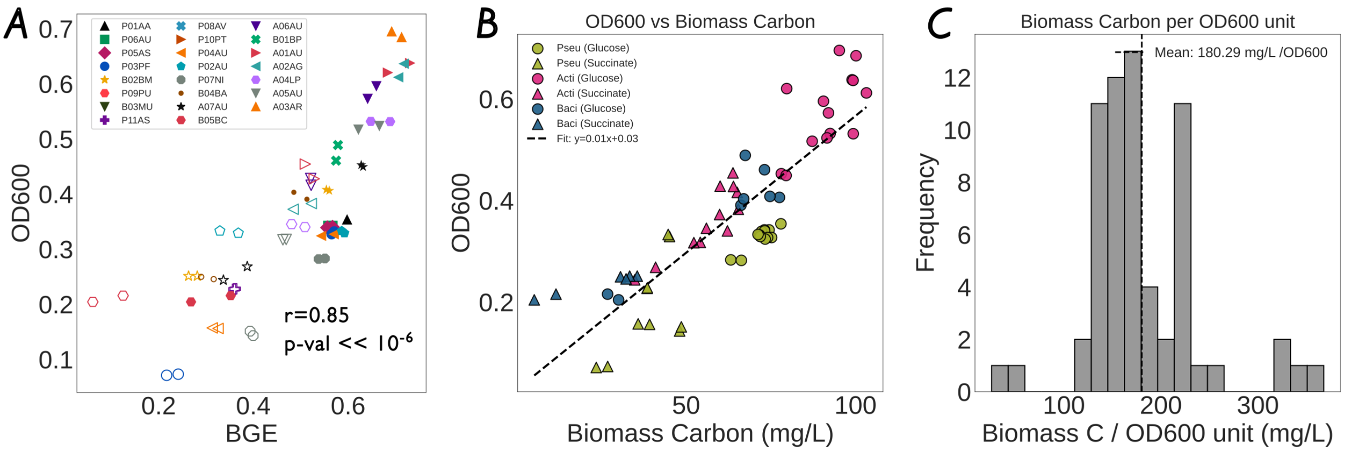


**Fig. S4.** $(\mathbf{A})$: BGE vs $OD_{600}$ at the end of the carbon quantification measurement. The OD yield is significantly correlated with BGE in our study. Spearman $r$: 0.85, p-value < $10^{-6}$. Each strain has a unique color and marker associated with it. Solid markers: glucose. Hollow markers: succinate.$(\mathbf{B})$:Biomass vs $OD_{600}$ yield. While there is a strong correlation between the two quantities, it is not a fixed quantity across resources and phyla. $(\mathbf{C})$ Distribution of Biomass Carbon/$OD_{600}$ unit from our measurements. (Two replicates per measurement. Both replicates shown in plots)

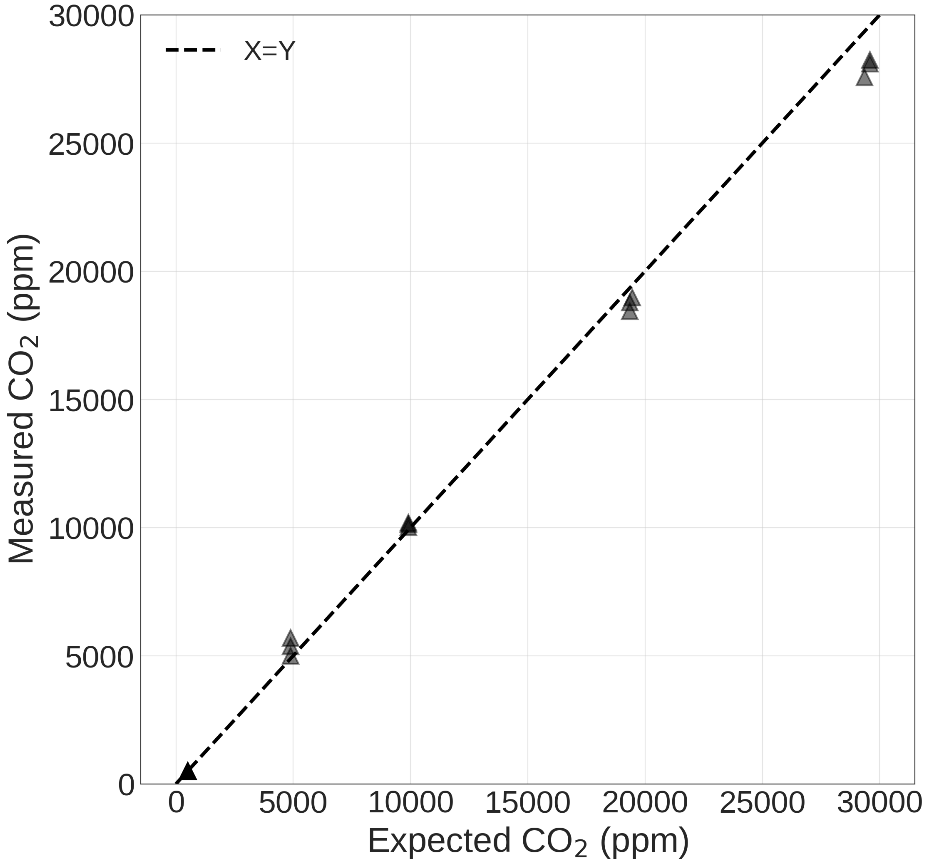


**Fig. S5.** SCD30 Calibration Curve: We tested the linear response of the SCD30 $CO_2$ sensors by adding increasing amounts of $NaHCO_3$ to dilute acetic acid; a reaction that releases a stoichiometrically known quantity of $CO_2$. Data shown is for a randomly selected sensor. Three replicates per calibration point.

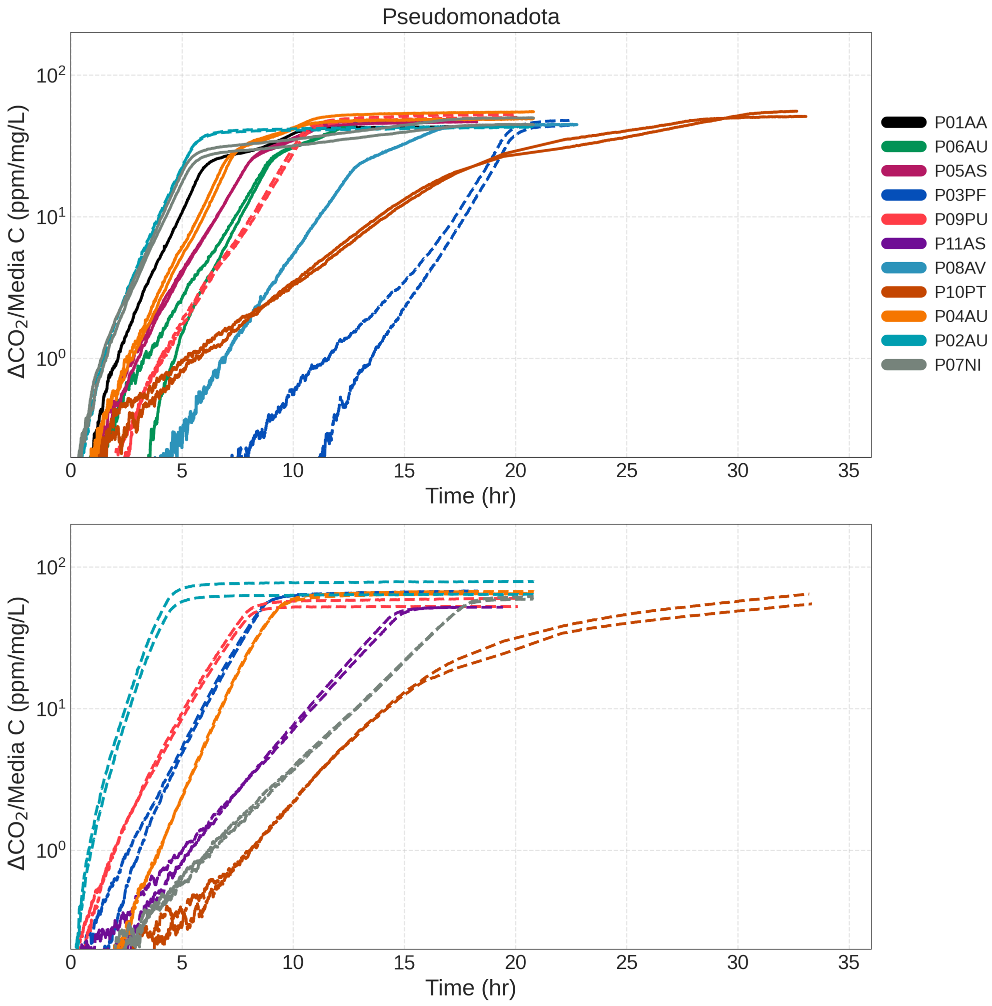


**Fig. S6.** $CO_2$ dynamics for all strains used in this study from the Pseudomonadota phyla. Y-axis: Respired $CO_2$ in ppm on a log axis, normalized to carbon concentration in growth media. X-axis time. Top panel: growth on glucose, bottom panel: growth on succinate. Two replicates per strain per resource condition. Trace colors are unique to each strain. Solid lines: biphasic respiration indicative of overflow metabolism, dotted lines: single-phase respiration indicative of no overflow metabolism.

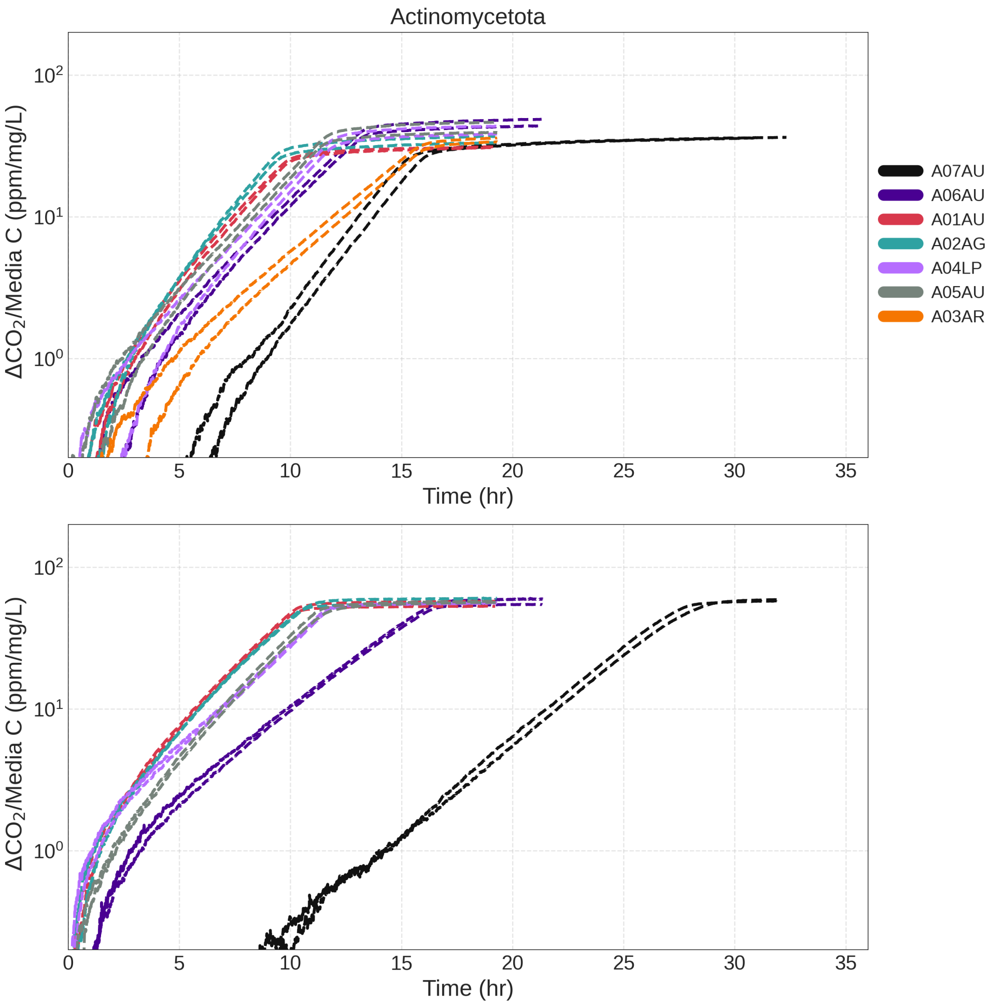


**Fig. S7.** $CO_2$ dynamics for all strains used in this study from the Actinomycetota phyla. Y-axis: Respired $CO_2$ in ppm on a log axis, normalized to carbon concentration in growth media. X-axis time. Top panel: growth on glucose, bottom panel: growth on succinate. Two replicates per strain per resource condition. Trace colors are unique to each strain. Solid lines: biphasic respiration indicative of overflow metabolism, dotted lines: single phase respiration indicative of no overflow metabolism.

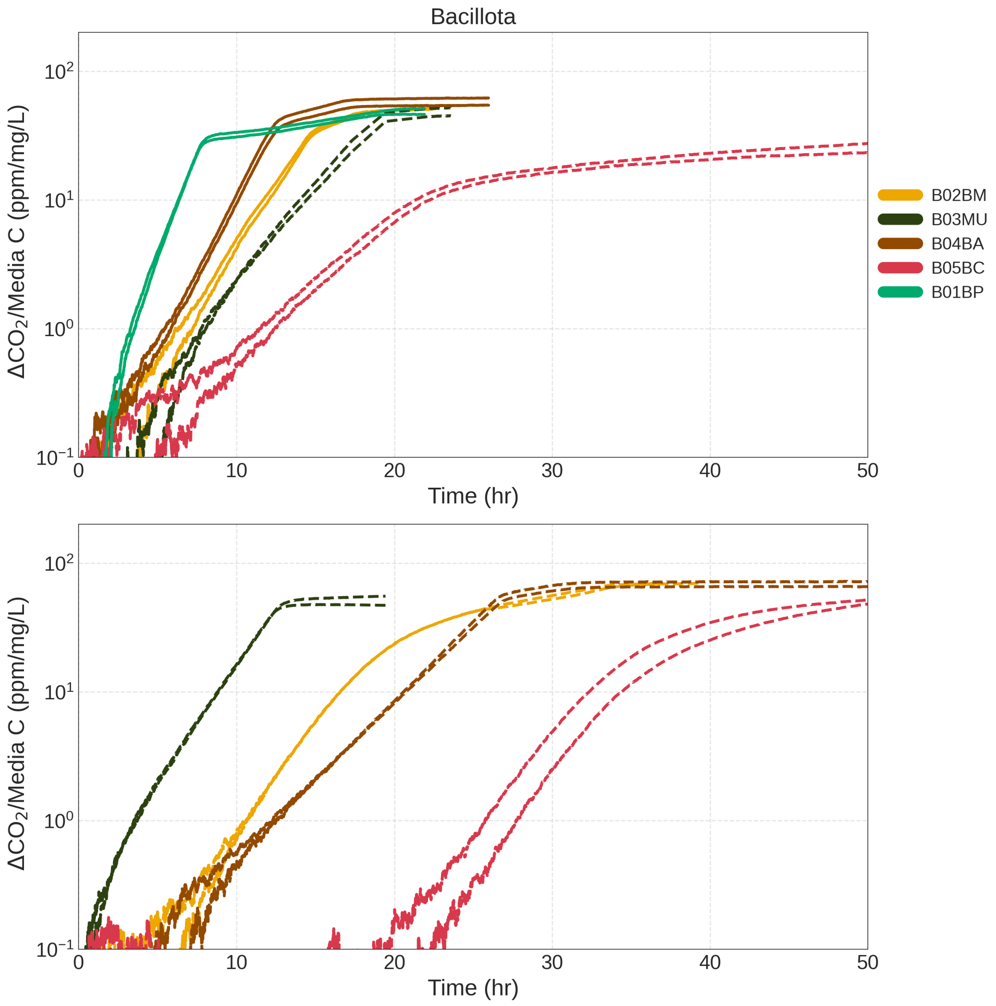


**Fig. S8.** $CO_2$ dynamics for all strains used in this study from the Bacillota phyla. Y-axis: Respired $CO_2$ in ppm on a log axis, normalized to carbon concentration in growth media. X-axis time. Top panel: growth on glucose, bottom panel: growth on succinate. Two replicates per strain per resource condition. Trace colors are unique to each strain. Solid lines: biphasic respiration indicative of overflow metabolism, dotted lines: single phase respiration indicative of no overflow metabolism.

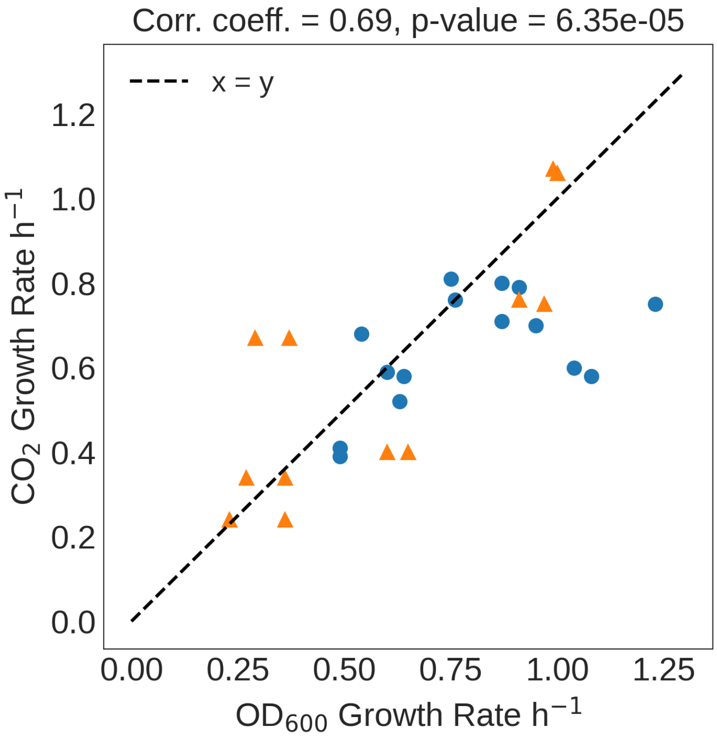


**Fig. S9.** Scatter plot of growth rate inferred by $OD_{600}$ measurements in a BMG platereader and with $CO_2$ measurements in respiration tubes. Circles: growth on glucose. Triangles: growth on succinate. Data plotted is mean across two replicates for each strain and resource combination. Only a subset of strains are shown here.

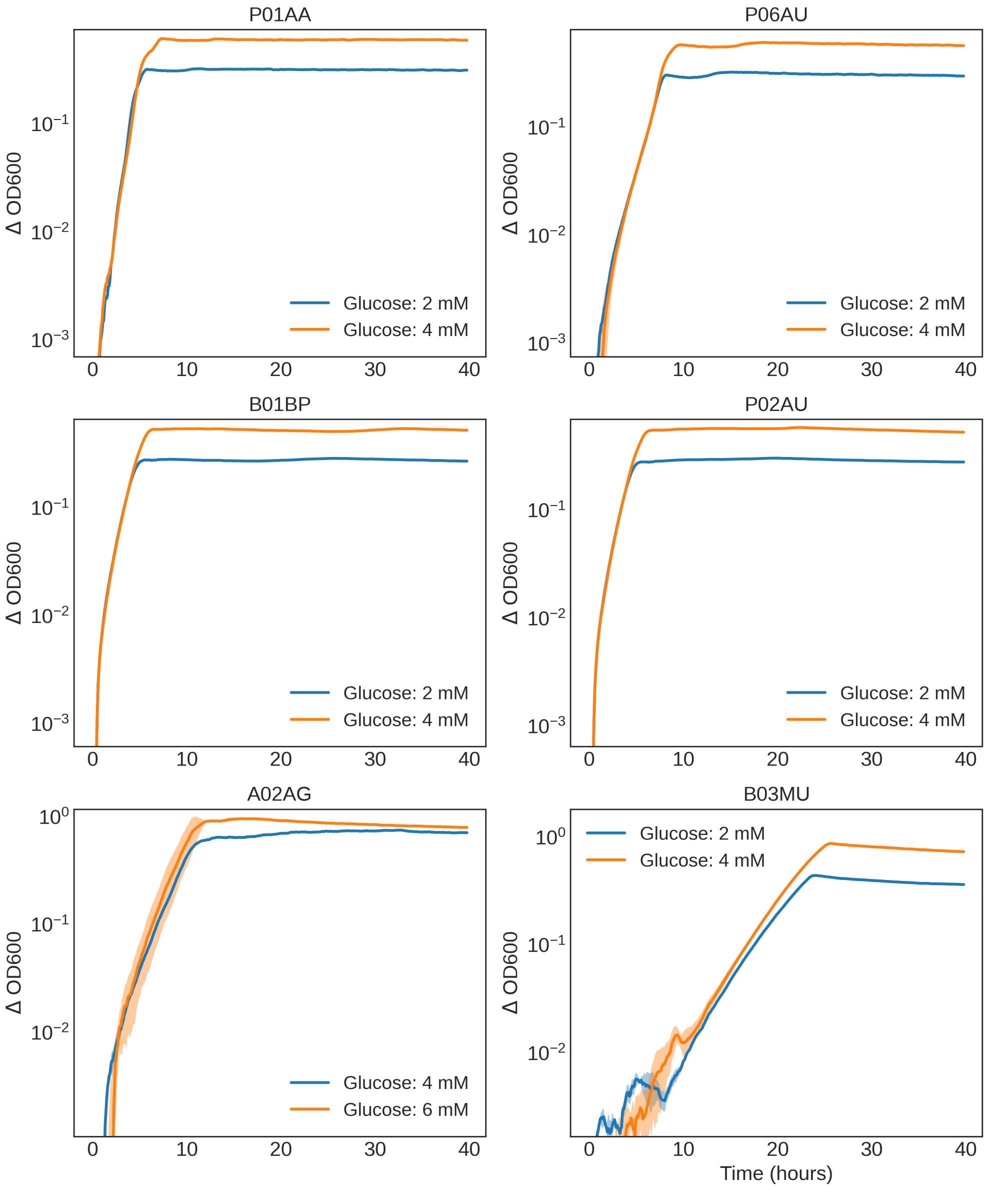


**Fig. S10.** $OD_{600}$ growth curves for six strains cultured in M9 medium supplemented with increasing concentrations of glucose (see legend for concentrations). Optical density was measured every 5 min, and values are shown on a log-transformed scale. Cultures for each strain and condition were inoculated at the same initial $OD_{600}$ following an identical preculturing protocol. Two biological replicates were measured for each strain and glucose concentration. Solid lines plot the mean of the replicates, and shaded envelopes plot the standard error. Growth rates for a given strain were consistent across glucose concentrations, whereas final biomass yield ($OD_{600}$) increased with higher glucose availability. This observation is consistent with our hypothesis that biomass growth on glucose is carbon-limited under the conditions of our experiment. Note that for strain A02AG, we chose the resource concentrations of 4 mM and 6 mM glucose, while for the other strains we chose 2 mM and 4 mM glucose.

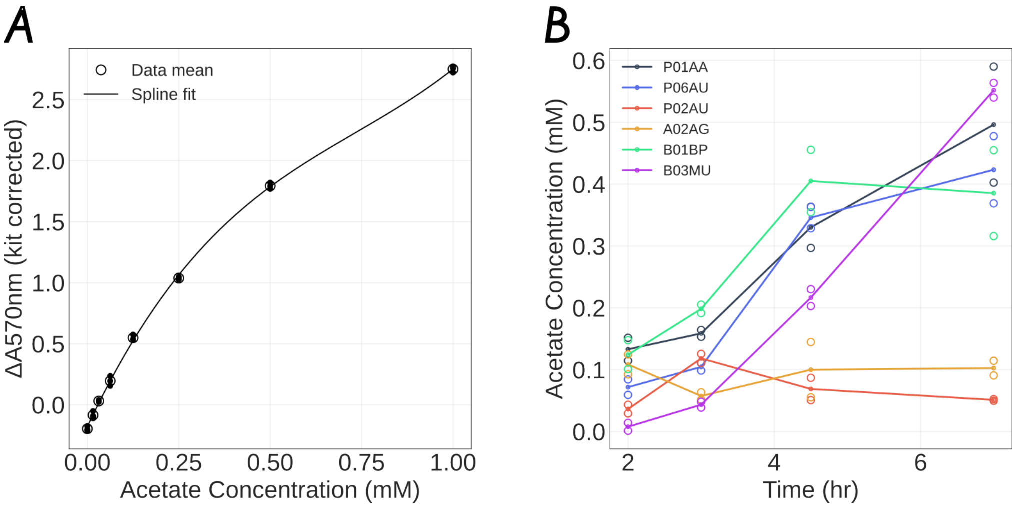


**Fig. S11.** Measurements of acetate production by strains growing on glucose. **(A)**: Calibration curve, corrected for background from a different kit. **(B)**: acetate production by chosen strains over time. (Two replicates per measurement. Both replicates shown in plots.)

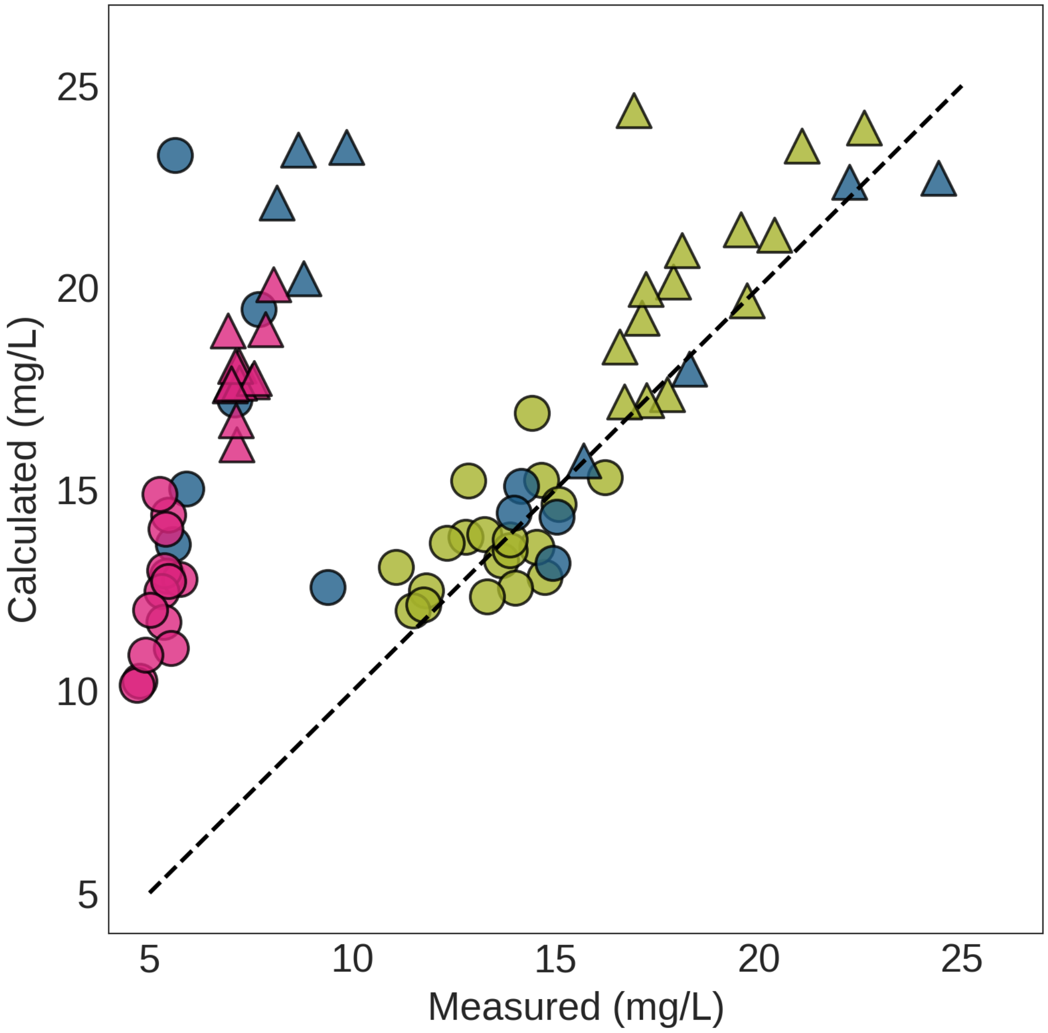


**Fig. S12.** Measured versus calculated concentration of dissolved carbonates in the culture at the end of the growth. The measurement is done using the TOC-L instrument. The calculated concentration is from the concentration of $CO_2$ in the headspace measured by the NDIR $CO_2$ sensors. Headspace $CO_2$ readings in ppm were converted to mg/L for comparison. (Two replicates per measurement. Both replicates shown in plot.)

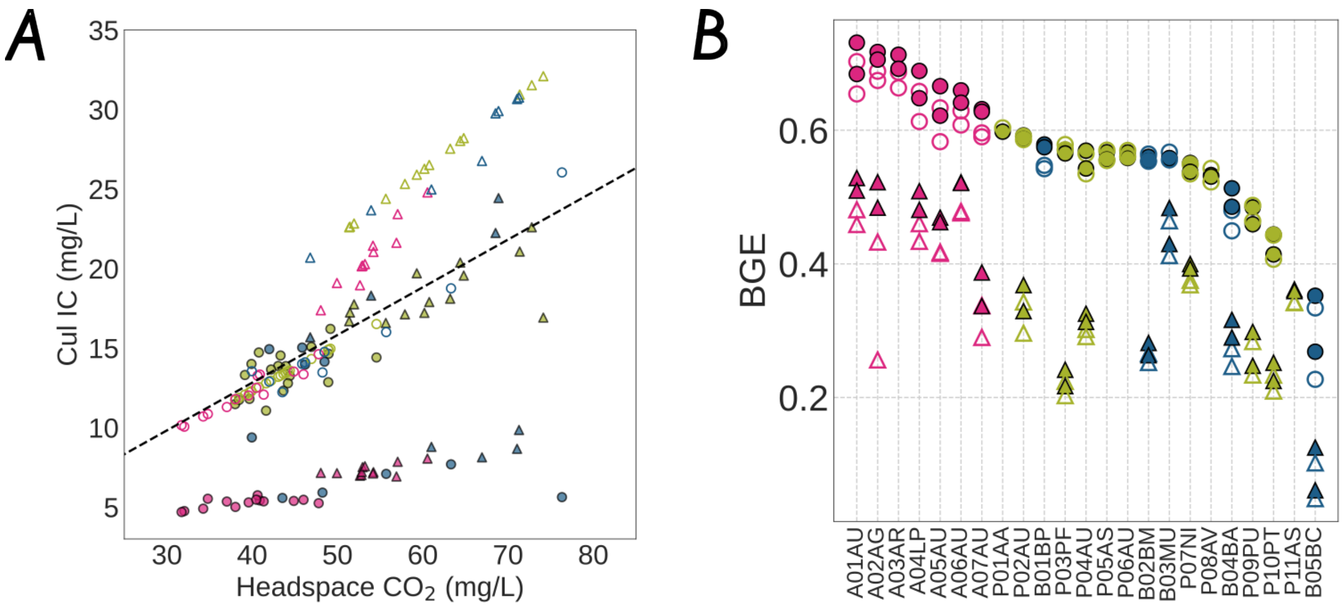


**Fig. S13.** **(A)**: Plotting measured headspace $CO_2$ (converted into mg/L of C dissolved in liquid culture volume) against dissolved inorganic carbon in the culture (labeled as CuI IC). Solid markers: measured $CO_2$ and measured CuI IC using the TOC-L instrument. Hollow markers: measured $CO_2$ and CuI IC calculated using Henry's law and the Henderson-Hasselbalch with pH measurements. There is a discrepancy between the measured and predicted CuI IC for some strains that we were not able to explain. **(B)**: BGE ranked across strains. Solid markers correspond to BGE computed from direct measurements. The total inorganic fraction needed to compute BGE consists of the headspace gaseous $CO_2$, dissolved $CO_2$, and inorganic carbon in carbonate form. We recalculated the BGE by using the headspace $CO_2$ to calculate the CuI IC component, plotted as hollow markers. As seen from the plot, accounting for the missing inorganic carbon has a small effect and in essence doesn't change our findings. (Two replicates per measurement. Both replicates shown in plots.)

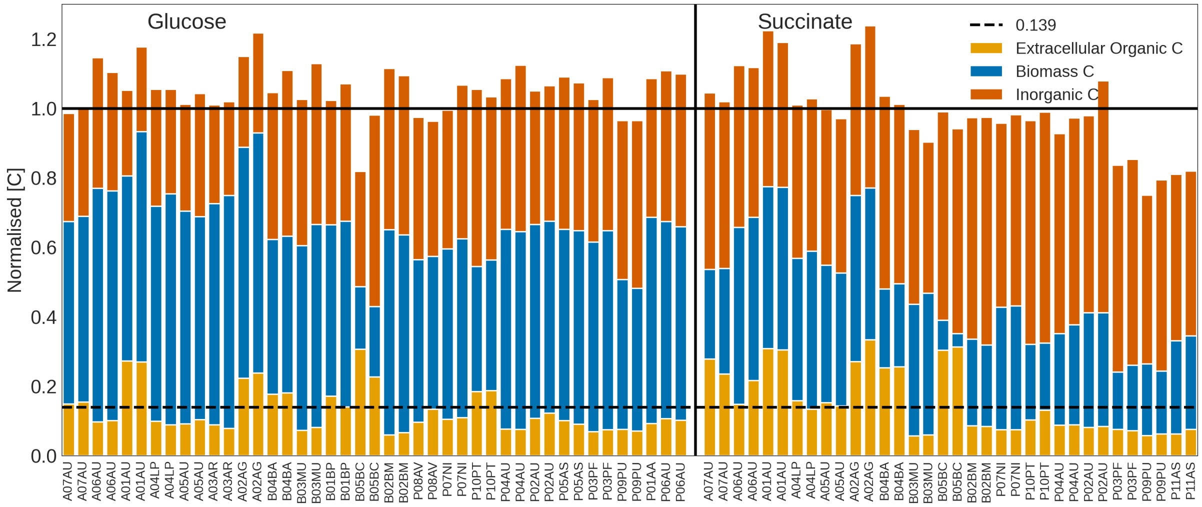


**Fig. S14.** All carbon pools measured directly using the TOC-L for strains used in this study. Y-axis: Measured carbon normalized to the provided media carbon. X-axis: Replicate measurements (2 replicates per strain, per resource). Strains are ordered by phyla. The left half of the plot corresponds to carbon pools for strains grown on glucose. The right half corresponds to succinate. We suspect the errors in the total carbon tally arise from error propagation from measurements across the TOC-L, the NDIR sensors, pipetting, as well as variance in the actual volume of the Borosil respiration tubes.

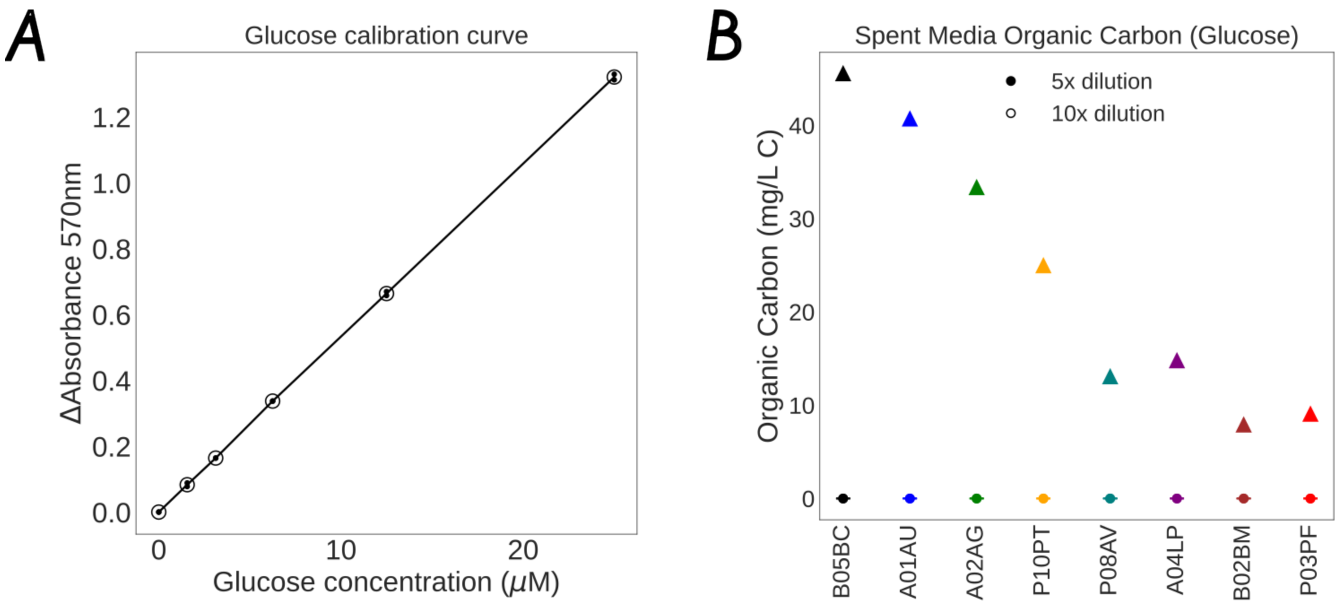


**Fig. S15.** Glucose assay of spent media. **(A)**: Calibration curve (2 replicates per concentration). **(B)**: Circular markers: Glucose concentration in spent media for eight strains grown on glucose, measured at 1/5X and 1/10X dilutions (2 replicates per dilution) and inferred from panel A; glucose concentrations were converted to mg/L C (6 carbons per glucose molecule) for direct comparison with TOC-L measurements in mg/L C. Triangular markers: Total organic carbon (TOC-L instrument). Across all strains, residual glucose is negligible relative to total organic carbon, indicating near-complete substrate consumption.

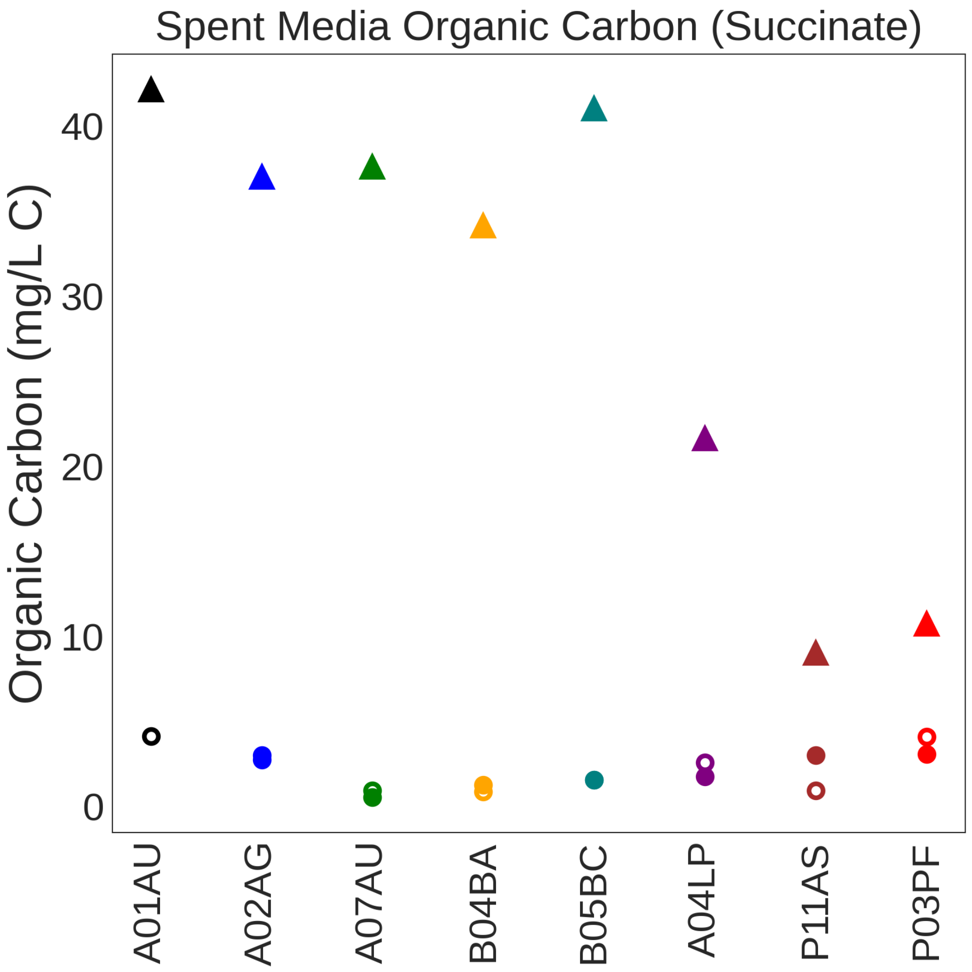


**Fig. S16.** Succinate concentration in spent media for eight strains grown on succinate, measured at 1/5X and 1/10X dilutions (2 replicates per dilution). Circular markers (left y-axis): inferred residual succinate after dilution correction, converted to mg/L C (4 carbons per succinate molecule) for direct comparison with TOC-L measurements. Triangular markers: Total organic carbon (TOC-L instrument). Residual succinate is negligible relative to total organic carbon, indicating near-complete substrate consumption. (The EnzyChrom ESNT-100 colorimetric assay requires no calibration curve.)

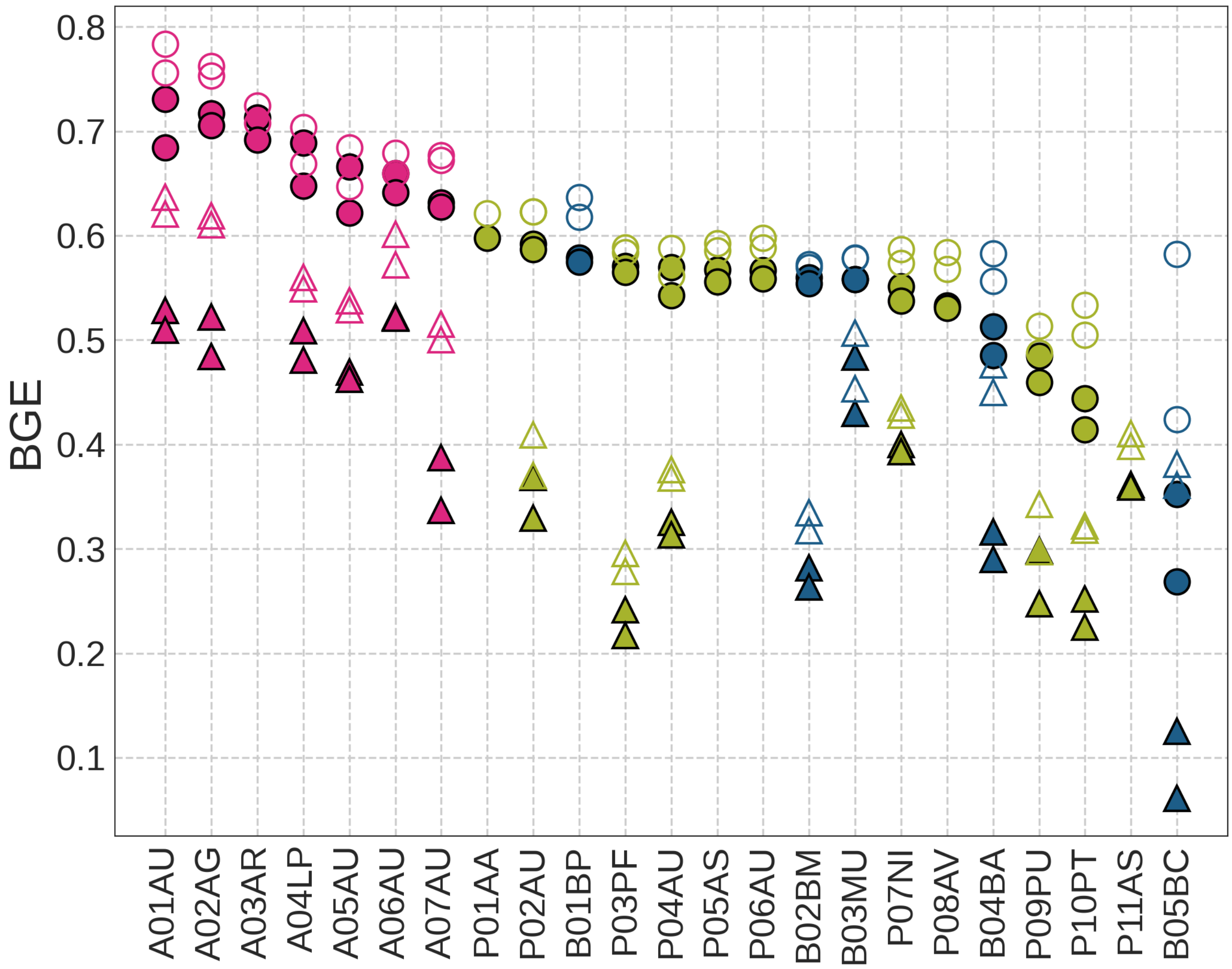


**Fig. S17.** BGE ranked across strains. Solid markers correspond to BGE reported in the main text that includes only biomass and inorganic carbon. Hollow markers correspond to BGE computed by assuming that the measured organic carbon in the spent media was also part of the biomass. Circular markers: glucose, triangles: succinate. Two replicates per strain and resource.

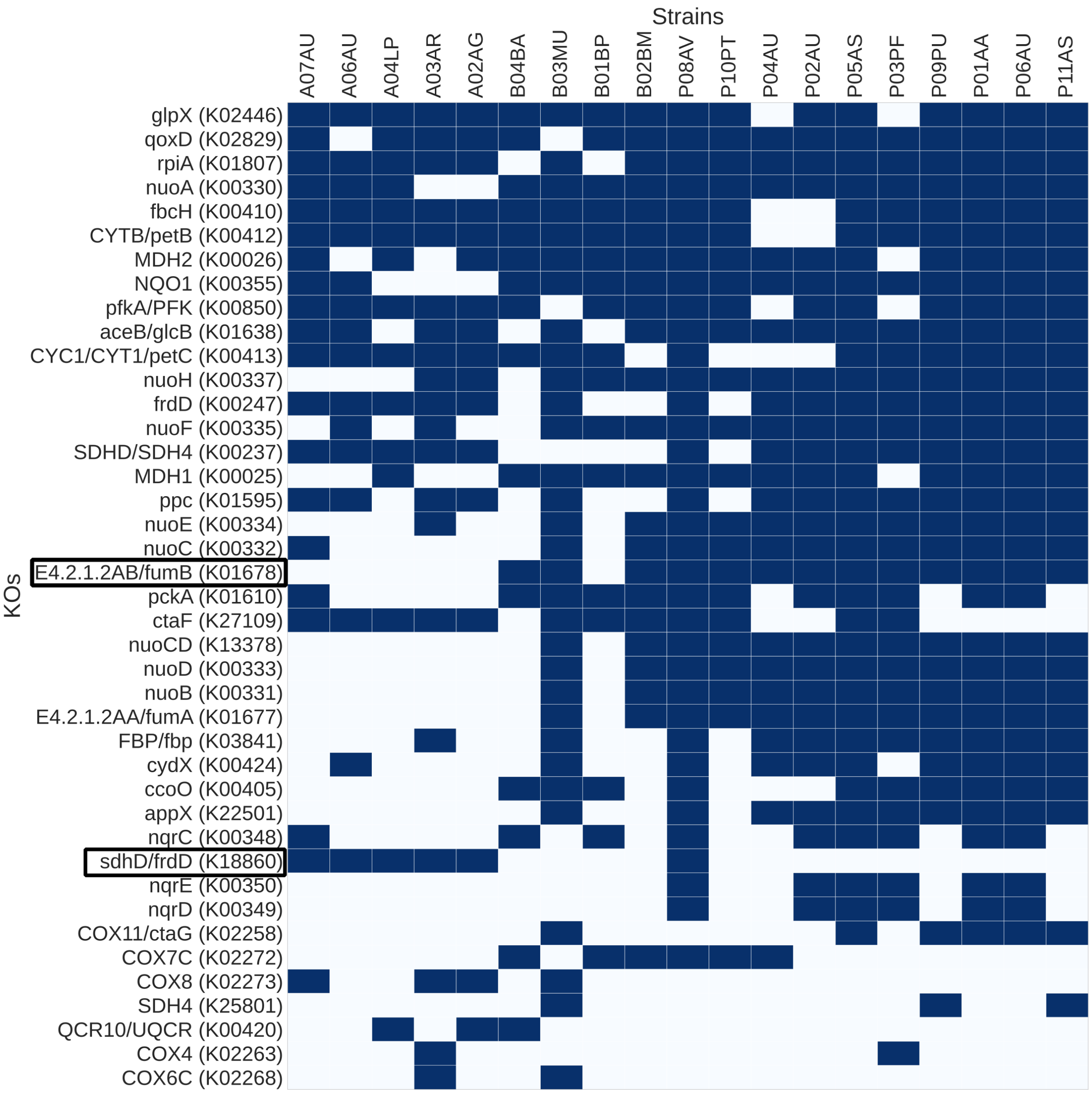


**Fig. S18.** Filtered gene-presence-absence table with 41 genes across the Electron Transport Chain and the TCA cycle. Blank cells indicate the gene (row) is missing in the strain (column). Blue cells indicate the strain has at least one copy of the gene. The 2 most predictive genes from our Random Forest analysis in the main text are marked along the y-axis: *fumB* and *sdhD/frdD*.

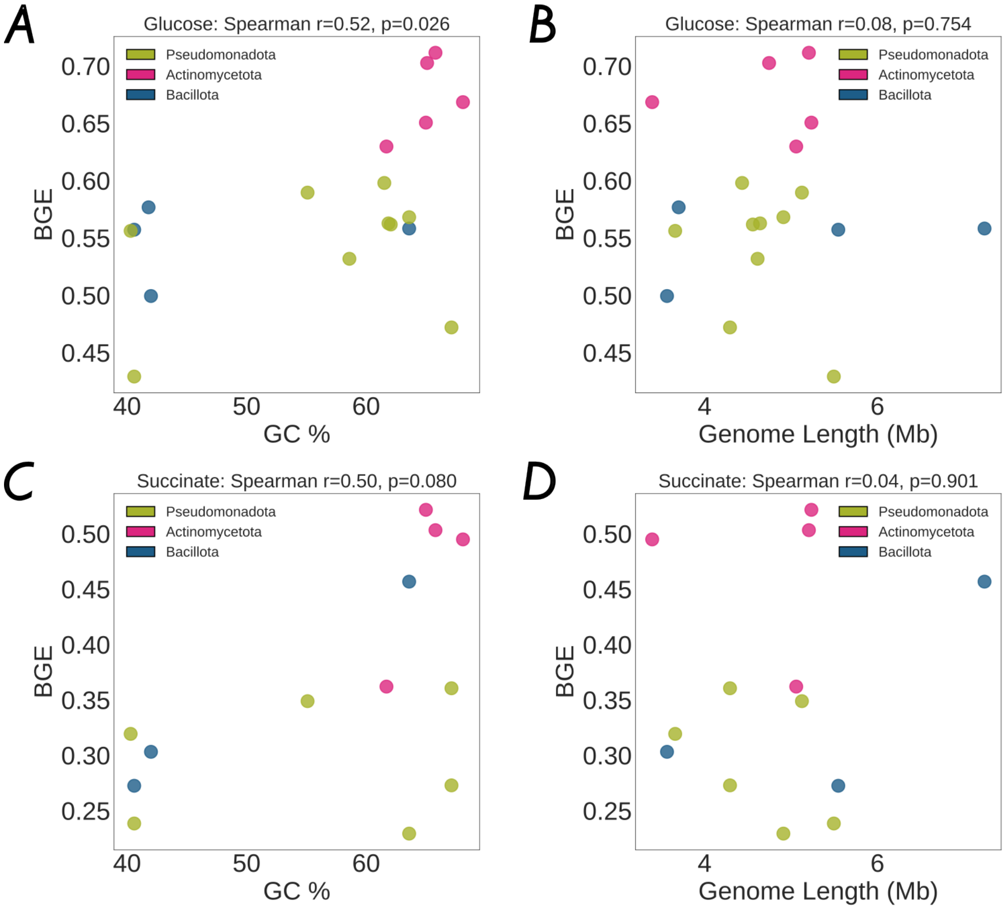


**Fig. S19.** Plots of BGE against genome statistics: length, GC content. We find that contrary to Saifuddin et al.(15), %GC content seems more positively correlated with BGE, and genome length has no correlation. **(A)** BGE on glucose vs %GC content of genome. **(B)** BGE on glucose vs genome length **(C)** BGE on succinate vs %GC content of genome. **(D)** BGE on succinate vs genome length

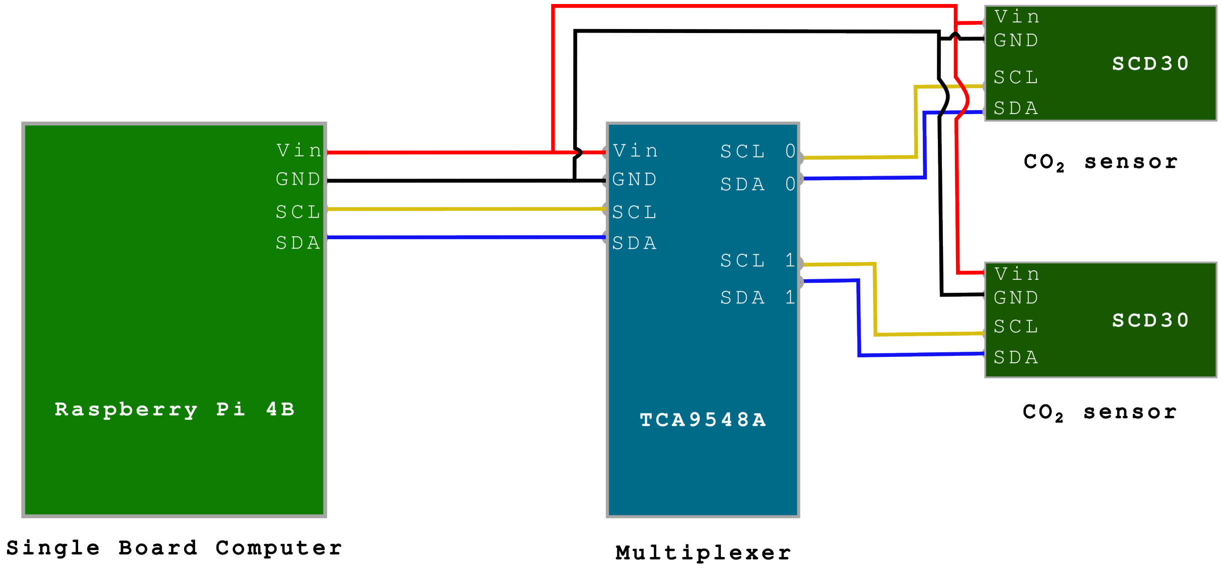


**Fig. S20.** Circuit diagram: Connecting a Raspberry Pi 4B to multiple SCD30 $CO_2$ sensors via a multiplexer module TCA9548A. Only two sensors are shown in this figure as examples, but the module can support up to 8.
